# Exploring Genetic Interactions with Rare Variants Reveals Gene Networks Susceptible to Complex Diseases

**DOI:** 10.1101/2024.12.17.628845

**Authors:** Hui Jiang, Bin Tang, Kun Li, Liubing Zhang, Junhao Liang, Clara Sze-Man Tang, Paul Kwong-Hang Tam, Binbin Wang, Youqiang Song, Qiang Wang, Mulin Jun Li, Hailiang Huang, Miaoxin Li

## Abstract

Genetic interactions play a crucial role in understanding the susceptibility and etiology of complex human diseases. However, existing methods for analyzing gene-gene interactions are often limited to common variants and may not capture the effects of rare variants. In this study, we propose a novel method called GGI-RUNNER that is specifically designed for analyzing gene-gene interactions of rare variants. GGI-RUNNER evaluates the enrichment of rare variant interaction burden in patients relative to baselines in the general population. The baselines are estimated for pairwise genes by a recursive truncated negative-binomial regression model on multiple genomic features available from public data. Intensive simulations demonstrate that GGI-RUNNER exhibits substantially higher statistical power than alternative epistasis tests and maintains reasonable type I error rates even in stratified populations. Applied to real data, GGI-RUNNER identified significant rare variant interactions associated with Alzheimer’s disease, Hirschsprung’s disease, Ulcerative colitis and Crohn’s disease. Furthermore, we found that these identified genes for each trait can form interconnected networks, which may provide valuable insights into the underlying molecular mechanisms.

## Introduction

Complex human diseases are often influenced by multiple genes and their interactions. Despite the advancements in genome-wide association studies (GWAS), the genetic basis of these diseases remains largely unexplained, leading to the issue of “missing heritability^1; 2^. Genetic interactions, or epistasis, which refers to the deviation from the additive effects of two or more genetic variants on the phenotype, are thought to be a potential source of unexplained genetic variation^3–5^. Epistasis has long been recognized as fundamentally important in the structure and evolution of genetic systems ^6^. The crucial role of epistasis in disease susceptibility and etiology has been revealed in complex human diseases, such as Alzheimer’s disease^7^, schizophrenia^8^, and inflammatory bowel disease^9^, and provided valuable insights into the complexities of their genetic architecture.

Methods for interaction analysis were originally explored at the single-nucleotide polymorphism (SNP) level, including parametric and non-parametric statistical methods, as well as machine learning and data mining techniques^10–15^. However, such methods designed to test for interactions within common variants are underpowered for analyzing rare variants (i.e., genetic variants with minor allele frequencies (MAF) <1%) unless sample sizes or effect sizes are extremely large. To increase power, collapsing strategies are extended to the study of gene-gene interactions (GGIs). The gene-based methods aggregate information across multiple rare variants inside a gene and collectively test interaction between variant pairs within two genes. In which odds ratio for FLR^16^, Pearson’s goodness-of-fit statistic for IGOF^17^, or multifactor dimensionality reduction (MDR) for GxGrare^18^, were employed to compare the difference in genotype distribution of a pair of genes between cases and controls. Then, a *P*-value for the tested gene pair was calculated based on specific distributions (i.e., chi-square distribution) ^16; 17^ or permutations^17; 18^. Despite the improved power compared to SNP-based epistasis tests, existing GGI tests for rare variants still exhibit limitations in genome-wide or exome-wide analysis due to insufficient sample sizes and excessive computing demand. Therefore, developing a powerful test will be needed for robust detection interactions between rare variants.

The development of reference population databases^19–21^ facilitated a series of databases for variant and gene interpretation and provided alternative approaches to map genotype-phenotype associations. Incorporating functional annotations, such as conservation scores^22; 23^ and predicted pathogenic scores^24–26^, has been widely accepted and successfully used to boost the power of rare variant association studies^27–30^. Recently, the RUNNER method^31^ further enhanced the power of rare variant association tests by accurately modeling the expected mutation burden across genes through regression of multiple genomic features obtained from publicly available databases. Compared to conventional rare variant methods like CMC^32^ and SKAT^28^, which rely on case-control differences in the number of rare mutation alleles, RUNNER demonstrates better performance in samples with small or medium sample sizes by comparing the observed rare mutation burden in patients against its baseline expectation in the population. Given the commonly encountered issue of inadequate sample sizes, the unique strategy of RUNNER, benefiting from leveraging external databases, shows great potential in enhancing statistical power and opening up new possibilities for gene-gene interaction analysis of rare variants.

Another major limitation in genome-wide analyses of gene interactions is the large number of interactions that need to be tested, which grows exponentially with the number of genes, specifically in an exhaustive search for pairwise interactions (*n*(*n* − 1)/2 tests for *n* genes). Such a search creates a heavy computational burden and statistical challenge for multiple testing corrections ^33^. Focusing on particular subsets of interactions with prior knowledge or functional relevance can effectively reduce the testing space. In this context, our previous work, the Digenic Interaction Effect Predictor (DIEP)^34^, provides a valuable resource for researchers to identify pathogenic gene pairs with potential digenic interaction effects. For instance, by considering gene pairs with a DIEP prediction score over 0.5 in human protein-coding genes (approximately 20,000 genes), the number of interactions can be reduced from approximately 400 million in an exhaustive search to nearly 4 million (1%). A more stringent threshold can further narrow the search space, thus substantially reducing the computing and multiple testing burden.

Here, we propose a novel gene-gene interaction test, GGI-RUNNER, as an extension of RUNNER to detect the joint interaction effects of multiple rare variants in a pair of genes on a quantitative trait. It uses a recursive truncated negative-binomial regression model to evaluate the enrichment of the rare variant interaction burden (RVIB) observed in patients relative to its expectation in the general population across pairwise genes, which is predicted based on multiple genomic features from external databases. In the present study, we performed extensive simulation studies to demonstrate that GGI-RUNNER maintains accurate false positive rates and achieves greater power in detecting rare variant interactions over a range of epistatic disease models. We also applied GGI-RUNNER to several real datasets of complex diseases: Alzheimer’s disease (AD), Hirschsprung’s disease (HSCR), Ulcerative colitis (UC) and Crohn’s disease (CD). We show that GGI-RUNNER is computationally efficient and identifies promising gene-gene interactions.

## Materials & Methods

### Notation and model

Suppose there are *K* affected individuals sequenced in the study. Consider two genes, *i* and *j*, and each gene is a collection of *V*_*i*_ and *V*_*j*_ rare variants (e.g., the variant with MAF < 1% in the reference population), respectively. Let *g*_*i,p,k*_ = 0, 1, or 2 represent the number of copies of the minor alleles for variant *p* in gene *i* carried by the *k*-th case subject (*p* ∈ *V*_*i*_, *k* = 1,2, …, *K*). A similar representation *g*_*j*,*q*,*k*_ is used for variant *q* in gene *j* (*q* ∈ *V*_*j*_). Here, we use rare variant interaction burden (RVIB) to measure the degree of rare variants with potential interaction effects on a pair of genes. A basic RVIB of the gene pair (*i*, *j*) observed in cases can be simply measured based on genotype functions as:

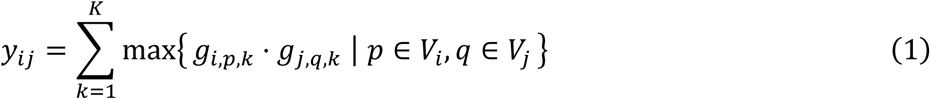

In the calculation, only pairs of variants with at least one copy of the minor allele present at both loci are considered potentially interacting. This assumption is consistent with the two-locus interaction model^35^. RVIB can be further improved by incorporating functional annotations of variants on a gene. Assume a variant *v* of gene *G* has a score, *s*_*v*,*G*_ ∈ [0,1], indicating its pathogenic potential. Given a score bin length *b* ∈ (0,1], *s*_*v*,*G*_ can be converted to an integer functional weight by the ceiling function as: *w*_*v*,*G*_ = [*s*_*v*,*G*_⁄*b*]. So, the functional weighted RVIB of the gene pair (*i*, *j*) is calculated as:

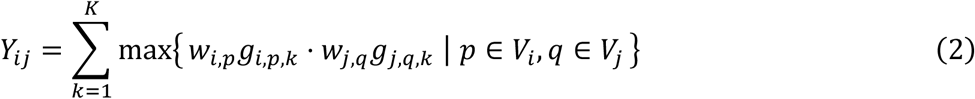

where *w*_*i*,*p*_ and *w*_*j*,*q*_ are the integer functional weight of rare variant *p* on gene *i* and variant *q* on gene *j*, respectively. In this study, functional weights were assigned based on integrated deleterious scores predicted using the logistic regression model embedded in the KGGSeq software ^36^. For downstream analyses, unless otherwise specified, we use the functional weighted *Y*_*ij*_ to calculate rare variant interaction burden for tested gene pairs. By analyzing the distribution characteristics in real data (Figure S1), we assume that the RVIB statistic *Y*_*ij*_ is asymptotically distributed as a truncated negative-binomial (TNB) distribution:

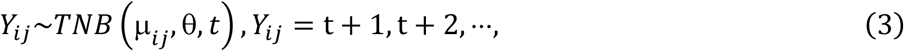

where μ*_ij_* is the expected baseline RVIB of the gene pair (*i*, *j*), and θ is a dispersion parameter. A notable distinction from the classical negative-binomial distribution is that a flexible data-dependent truncation point *t* is introduced in the TNB distribution to restrict the range of values for *Y*_*ij*_. Details about the TNB distribution were described in Supplemental Methods.

By default, GGI-RUNNER uses prediction scores from DIEP^34^ to narrow all pairwise genes down to a subset of pairs with potential pathogenic interaction effects. The DIEP scores were precalculated for all protein-coding gene pairs across the human genome, and 3,920,174 gene pairs were predicted with a DIEP score over 0.5. The distribution of DIEP scores ([0,1]) can be seen in Figure S2.

### GGI-RUNNER estimates the expected RVIB based on multiple genomic features

Based on the TNB distribution, GGI-RUNNER constructs a generalized linear regression model to estimate the expected RVIB in the general population to test the enrichment of rare variant interactions at gene pairs in patients. In the present study, six genomic features that can potentially impact the gene’s number of rare mutation alleles were considered in the regression model. For a single gene, these features include the coding region (CDS) length (*X*_1_), the accumulated MAF (*X*_2_), the product of CDS length and MAF (*X*_3_), the observed/expected ratios for missense (oe_mis, *X*_4_) and loss-of-function variations (oe_lof, *X*_5_), and the GC content (*X*_6_). Values for these six features can be obtained or calculated from the gnomAD database^19^, a detailed calculation for the *X*_2_ is given in Supplemental Methods. When analyzing a gene pair (*i*, *j*), the value for each feature is calculated as the product of the corresponding feature values of gene *i* and gene *j*. Additionally, a similar rare variant interaction burden of the gene pair (*i*, *j*) observed in controls can be calculated and included in the regression model as the seventh predictor *X*_7,*ij*_. However, if the number of control subjects is limited (i.e., < 50) in the studied sample, *X*_7,*ij*_ will not be considered in the regression model, so we added an asterisk to this term in the equation below. Under a link function, the expected RVIB of gene pair (*i*, *j*) in the studied sample can be estimated as:

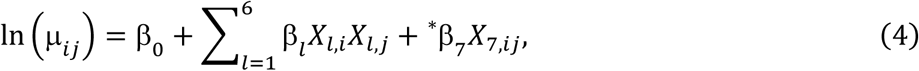

where β_0_ is an intercept, and β_1_ to β_7_ are the regression coefficients for *X*_1_ to *X*_7_, respectively. Since the vast majority of rare variant interactions between genes are not associated with disease status, we can simply let μ_*ij*_ = *Y*_*ij*_ in estimating the regression coefficients. Given a specific score bin length *b* for integerizing the function weight and a truncation point *t*, then the regression coefficients (β_0,_ β_1_, …, β_6_, β_7_^∗^) and dispersion parameter (θ) of the TNB distribution model can be estimated by maximum likelihood with a Quasi-Newton method. With the fitted coefficients, the deviance residuals are calculated and standardized as *d́*_*ij*_ (see details in the Supplemental Methods). A large *d́*_*ij*_ indicates the gene pair (*i*, *j*)’s observed RVIB in patients is greater than its expected RVIB in the general population. *d́*_*ij*_ can be approximated using a standard normal distribution N(0,1), and its *P*-value can be obtained analytically without premutation.

In practice, we do not know which gene pairs are not associated with disease status (the background gene pairs), so we propose employing a recursive regression procedure. First, we fitted the model with all the gene pairs and identified the gene pairs with significant and suggestively significant *P*-values (e.g., Benjamin– Hochberg false discovery rate (BHFDR) *q*-value ≤ 0.1). We then removed these gene pairs and re-fitted the model with the remaining gene pairs. This process was repeated until no gene pair had a large deviance residual or the significant gene pairs became stable. Then, the resulting set of gene pairs formed the background gene pairs. During each iteration, the regression coefficients were re-estimated using maximum likelihood with a Quasi-Newton method under the TNB distribution. Then, we used the estimated coefficients to calculate *P*-values for all the gene pairs, which were used to identify gene pairs for removal in the next iteration. This procedure allowed us to model the background gene pairs while minimizing the influence of real disease-associated gene pairs with excessive rare variant interaction burden on the coefficient’s estimation.

The optimal values for score bin length *b* and truncation point *t* were explored under a grid-search procedure. We tested values of *b* ranging from 0.025 to 0.5 with an interval of 0.025, and values of *t* ranging from 0 to 2 with a break of 1. The optimal combination of *b* and *t* was determined as the value minimizes the mean log fold change (MLFC)^37^, which measures the departure of *P*-values of background gene pairs from the uniform distribution, and maximizes the number of significant gene pairs, according to a balanced ranking. To prevent inflation of type I error rate, the recursive procedure under the specific combination of *b* and *t* will be stopped if the number of gene pairs used for regression fitting is less than 5000. Finally, GGI-RUNNER calculated *P*-values for all tested gene pairs based on the stable regression model using the optimal combination of *b* and *t*, and provided the results to the users.

### Experiments on simulation data

We used a semi-simulation procedure to test the performance of GGI-RUNNER in binary traits compared with PLINK^10^ and GxGrare^18^. PLINK is a widely used tool for testing SNP×SNP pairwise interactions on a genome-wide scale. PLINK provides two kinds of epistasis tests, which are implemented in the ‘--epistasis’ command and the ‘--fast-epistasis’ command. The ‘--epistasis’ option in PLINK (PLINK-epi) uses a logistic regression test for interaction that assumes an allelic model both for main effects and interactions. The ‘--fast-epistasis’ option in PLINK (PLINK-fast) performs a faster test based on a Z-score for the difference in SNP×SNP association (odds ratio) between cases and controls. GxGrare is a gene-gene interaction method designed for rare variants by performing the multifactor dimensionality reduction (MDR) analysis for the collapsed rare variants. GxGrare provides six permuted *P*-values for each pair of genes calculated based on different collapsing methods (MAF-based collapsing, functional region-based collapsing and effect-based collapsing) and evaluation measures (information gain and balanced accuracy). In our manuscript, we adopt the *P*-values of information gain using MAF-based collapsing for comparison according to the recommendation of GxGrare. The *P*-value is calculated by 1,000,000 permutations.

We first investigated the properties of our test through simulations on real exome data from the Singapore 10K Genome Project (SG10K)^38^. The SG10K database consists of whole-genome sequencing data of 4810 Singaporeans. After applying the KING method^39^ to remove relatedness up to the third degree (please refer to the Supplemental Methods for more details), a subset of SG10K that contains 4016 unrelated individuals was used for the simulation. To evaluate the performance of GGI-RUNNER on stratified populations, five ancestry panels of the 1000 Genomes Project (1KGP)^21^ were used for the simulation, including African (AFR, *N* = 661), American (AMR, *N* = 347), East Asian (EAS, *N* = 504), European (EUR, *N* = 503) and South Asian (SAS, *N* = 489).

#### Type I error simulations

For type I error simulations, we analyzed under the null hypothesis that all tested gene-pair interactions are not associated with disease risk. We first considered four sets of sample sizes, which we set to 1000, 2000, 3000 and 4000. Unrelated subjects were randomly selected from the SG10K subset to form a new sample comprising equal numbers of pseudo-cases and controls. Only rare variants with an MAF of less than 1% in the East Asian panel of gnomAD exomes and gnomAD genomes were included in the analysis. GGI-RUNNER analysis was performed on gene pairs with DIEP scores over 0.9. Average type I error rates were calculated based on 100 simulated replicates of samples at the significance level of α= 10^−3^, 10^−4^, 10^−5^. The distribution of *P*-values was assessed using the quantile-quantile (QQ) plot of −log_10_*P* under the uniform distribution U(0,1), and the MLFC was used to measure the departure of *P*-values from the uniform distribution. Additionally, we examined the impact of the MAF cutoff on the type I error rates. GGI-RUNNER was applied using a range of MAF cutoffs (i.e., 0.1%, 0.5%, 1% and 5%) with a fixed sample size of 4000.

To evaluate the effects of population stratification on type I error rates, we utilized subjects from the 1KGP dataset to create ancestral mixed samples. In each sample, two-thirds of individuals were randomly selected from a specific ancestry panel to form the pseudo-cases. The remaining one-third of individuals from the same panel and an equal number of individuals randomly drawn from another ancestry panel constituted the controls. Overall, we generated 20 samples by varying the panel combinations. GGI-RUNNER was performed on rare variants with an MAF of less than 1% in the gnomAD panel that matched the ancestry of the cases.

#### Power simulations

For empirical power simulations, we worked under the alternative hypothesis by inserting the causal mutations into the real genome. Unrelated subjects were randomly drawn from the SK10K subset and assigned as pseudo-cases or controls. The gene pair consisting of *PARK7* and *PINK1* was designated as the putative risk gene pair. To facilitate comparison with alternative methods, we assumed that the presence of only one pair of rare variants associated with susceptibility within the target gene pair of *PARK7*-*PINK1* (further details can be found in Table S1). Genotypes of all variants within these two target gene regions were pre-removed.

The simulation design used three built-in two-locus interaction models to generate epistasis scenarios, including the threshold model^35^, the multiplicative model^35^, and the classic epistasis model^11^. Table 1 shows these models in terms of the odds of diseases for each combination of genotypes at two loci A (disease risk allele a) and B (disease risk allele b). The symbols *α* and *θ* in the table represent the baseline risk and effect size, respectively. For a specific model, the *α* and θ can be calculated based on a pre-specified disease prevalence Pr(D) and a genotype odds ratio (OR) for an assumed minor allele frequency (MAF) of disease-associated variants^40^. More details of these models are provided in Supplemental Methods. In the simulation, the prevalence Pr(D) was fixed as 1% in all epistasis scenarios. For each model, we generate genotype data of assumed risk variants by setting the MAFs as 1% for each locus, and varying the OR (i.e., OR=5, 10, 15 and 20) and the sample size *N* (i.e., *N* =1000, 2000, 3000 and 4000) under a balanced case-control design. To evaluate the effects of MAF on power, we simulated genotypes for the disease variants under a range of MAFs (i.e., MAF=0.1%, 0.25%, 0.5%, 1% and 2% for each locus) with a fixed OR of 20, and simulated the disease status for 2000 cases and 2000 controls for each model. For each setting, we generated 100 datasets for each model. The power was estimated as the proportion of the simulated casual gene pair detected as a significant signal among 100 replicates.

**Table 1.** Odds tables for the three epistasis disease models.

| Model 1: The threshold model |  |  |  |
| --- | --- | --- | --- |
| Genotype | AA | Aa | aa |
| BB | $\alpha$ | $\alpha$ | $\alpha$ |
| Bb | $\alpha$ | $\alpha (1 + \theta)$ | $\alpha (1 + \theta)$ |
| bb | $\alpha$ | $\alpha (1 + \theta)$ | $\alpha (1 + \theta)$ |
| Model 2: The multiplicative model |  |  |  |
| Genotype | AA | Aa | aa |
| BB | $\alpha$ | $\alpha$ | $\alpha$ |
| Bb | $\alpha$ | $\alpha (1 + \theta)$ | $\alpha (1 + \theta)^2$ |
| bb | $\alpha$ | $\alpha (1 + \theta)^2$ | $\alpha (1 + \theta)^4$ |
| Model 3: Classic epistasis model |  |  |  |
| Genotype | AA | Aa | aa |
| BB | $\alpha$ | $\alpha$ | $\alpha (1 + 4\theta)$ |
| Bb | $\alpha$ | $\alpha (1 + 2\theta)$ | $\alpha$ |
| bb | $\alpha (1 + 4\theta)$ | $\alpha$ | $\alpha$ |

### Application to real data

We then applied GGI-RUNNER to four real high-throughput sequencing (HTS) datasets, including two local complex disease datasets: Alzheimer’s disease (AD)^41^ and Hirschsprung’s disease (HSCR)^42^, and two inflammatory bowel disease datasets from the UK Biobank whole-exome sequencing (WES) data^43^: Ulcerative colitis (UC) and Crohn’s disease (CD). Sample information of the four datasets is summarized in Table S2. All sequenced samples in AD and HSCR studies were obtained with Institutional Review Board approval in Hong Kong or mainland China. Human genetics data from the UK Biobank was conducted under applications No.15422 and No.86920.

All analyses for detecting susceptibility gene-gene interactions with rare variants from the HTS data were conducted in our KGGSeq platform (V1.2, http://pmglab.top/kggseq/). Quality control (QC) was performed in the following steps. We first removed the variants with the Hardy-Weinberg equilibrium *P*-value ≤ 1 × 10^−5^ and alleles of more than four types. Second, genotypes with read depth less than eight or a genotyping quality score (Phred Quality Score) < 20 were changed to a no-call. Finally, variants with more than 20% missing genotypes were excluded. After QC, we performed GGI-RUNNER analysis on rare non-synonymous variants (missense, start-loss, stop-loss, stop-gain, splicing, frameshift, and non-frameshift variants that were annotated by RefGene^44^ and GENCODE^45^) for protein-coding gene pairs with DIEP scores over 0.8. Table S2 also provides information on the number of rare variants and the number of gene pairs tested in each dataset.

In the discovery phase, STRING database^46^ (https://string-db.org/) was queried to identify published information on protein-protein interactions (PPIs). PPIs of five categories (including experimentally determined, curated databases, gene co-occurrence, co-expression and protein-homology) with confidence levels exceeding 0.5 were used for visualization. Gene-gene interactions were visualized using Cytoscape software^47^. To identify the biological functions of the genes identified by GGI-RUNNER, Gene Ontology (GO)^48^ analysis was executed on clusterProfiler (version 4.2.2) package^49^ in R software. The enrichment significance *P*-values were generated based on hypergeometric tests.

## Results

### Overview of methods

GGI-RUNNER is an innovative statistical method for analyzing gene-gene interactions of rare variants. It evaluates whether rare variant interactions between pairwise genes are significantly enriched in disease risk by comparing the observed rare variant interaction burden (RVIB, see details in Methods) in patients to its baseline expectation in the general population. There are three main steps in the GGI-RUNNER framework: (1) calculating the rare variant interaction burden in cases for pre-tested gene pairs and preparing predictor values based on public annotation databases; (2) recursively fitting the truncated negative-binomial regression model on background gene pairs, and (3) testing for significance of the deviation from the baseline rare variant interaction burden for all tested gene pairs based on the fitted regression model (Figure 1).

**Figure 1.**
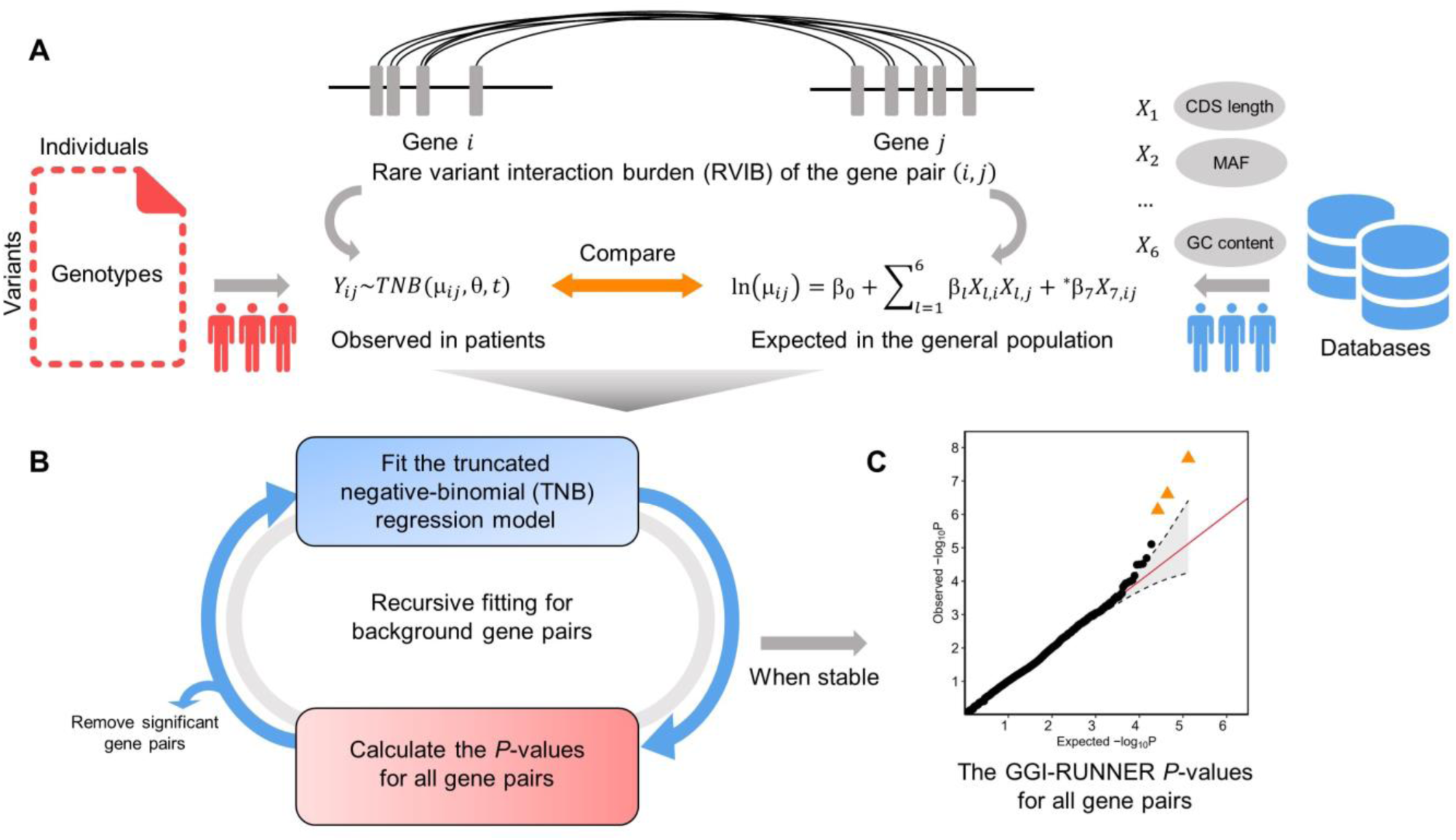
GGI-RUNNER workflow. **(A)** The GGI-RUNNER test evaluates the enrichment of rare variant interaction burden (RVIB) in patients compared to its baseline expectation in the general population. For each pre-tested gene pair, GGI-RUNNER calculates the observed RVIB in cases based on the user-input genotype data (left part), and estimates the expected RVIB by regressing multiple genomic features from external public data (right part). **(B)** Recursive fitting the truncated negative-binomial regression model to obtain a stable model constructed on background gene pairs. **(C)** The GGI-RUNNER *P*-values for all tested gene pairs are calculated and reported.

A key technical challenge for GGI-RUNNER is accurately estimating the baseline rare variant interaction burden for gene pairs. To address this challenge, we utilized a truncated negative-binomial regression model to estimate the baseline expectation based on multiple genomic features from public annotation databases (Figure 1A and Methods). This model offers advantages over the standard negative-binomial model by incorporating a data-dependent truncation point *t*, which accommodates the inflated number of gene pairs with zero or low rare variant interaction burden. Additionally, GGI-RUNNER employs a recursive regression procedure, similar to outlier removal in regression analysis, to model background gene pairs and generate baseline expectations under the null hypothesis (Figure 1B). This approach helps establish a reliable baseline expectation for rare variant interaction burden. To assess the significance of gene-gene interactions, GGI-RUNNER compares the rare variant interaction burden observed in patients with its estimated baseline expectation and calculates a corresponding *P*-value for each tested gene pair (Figures 1A and 1C). Notably, since this comparison is essentially only within the cases, the presence of population structure in the control subjects has minimal influence on the statistical tests of GGI-RUNNER. This design mitigates potential biases associated with confounding factors between cases and controls and enables more accurate identification of gene pairs with true significant interactions.

### Type I error simulations

The empirical type I error rates estimated for GGI-RUNNER are presented in Table 2 at α = 10^−3^, 10^−4^, 10^−5^ with four different sample sizes: 1000, 2000, 3000 and 4000 under a balanced case-control design. The results show that the false positive rate is protected across considered scenarios at all significance levels. The distribution of *P*-values for tested gene pairs is close to the uniform distribution U[0,1] (Figure S3), even at small sample sizes. The departure of *P*-values from the uniform distribution is measured by the MLFC (mean log fold change). As expected, we observed that the MLFCs for GGI-RUNNER were stable and moderate (< 0.2) in the analysis across all four different sample sizes and four MAF thresholds (Table 2).

**Table 2.**
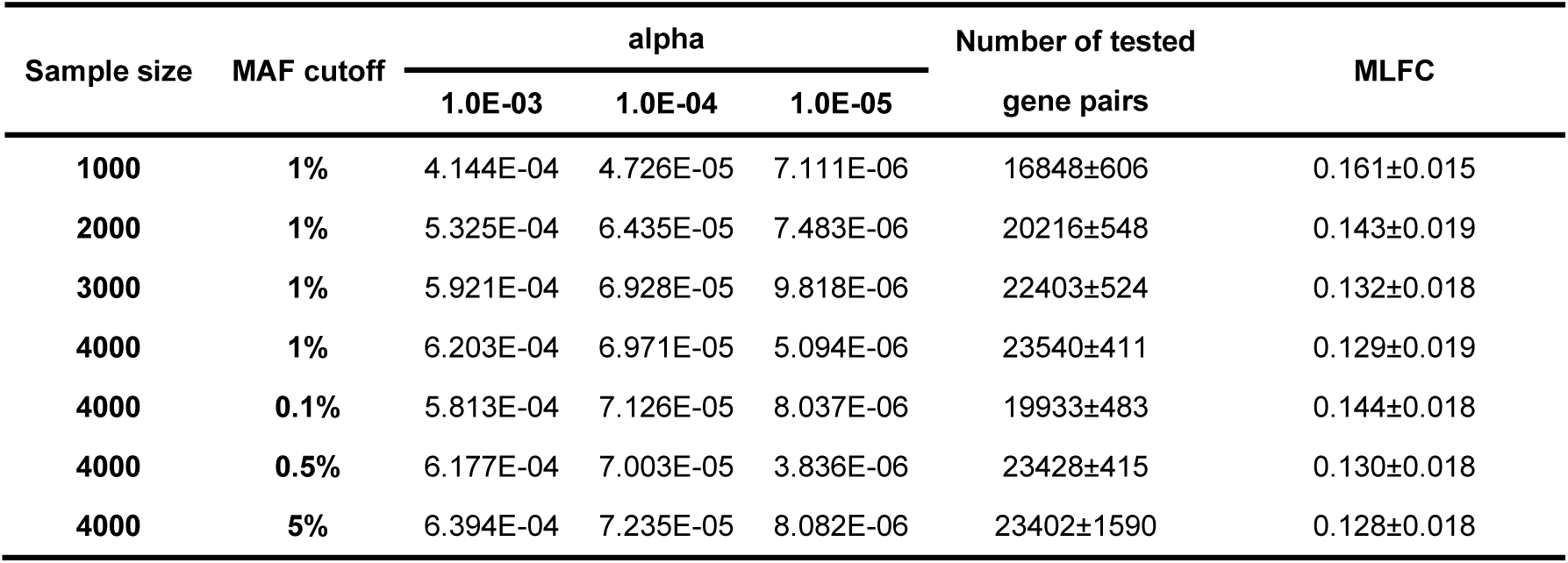
Type I error rates of GGI-RUNNER for testing interaction between two genes involving rare variants.

GGI-RUNNER was applied to samples with different sample sizes, 1000, 2000, 3000 and 4000, and tested gene-gene interactions of rare variants under different MAF cutoffs, 0.1%, 0.5%, 1% and 5%. The semi-simulation procedure produced the samples based on whole-genome sequencing data from SG10K unrelated subjects under a balanced case-control design. Each setting repeats 100 times. MLFC: the mean log fold change.

We then investigated the impact of population stratification by performing GGI-RUNNER on 20 ancestral mixed samples simulated based on 1KGP data. The results showed no inflation of spurious association even when 50% population stratification was present in controls, as evidenced by the QQ plots in Figure S4. The inflation index, MLFC, remained as low as observed in the above ethnically homogeneous populations (< 0.2) in all 20 samples. Likewise, most samples had no systematic type I error inflation (Table S3). While in some samples, a few gene pairs reached study-wide significance (BHFDR *q*-value < 0.05), as shown in the QQ plots (Figure S4). However, it should be noted that they were not randomly occurring as they could be detected in tests with different controls, suggesting that these gene pairs may be sample- or ethnic-specific. Therefore, GGI-RUNNER is robust to population stratification and can effectively control for its impact on rare variant interaction analysis.

### Empirical power simulations

Next, we evaluated the power of GGI-RUNNER for testing the interaction between two genes in a variety of configurations and compared the results with three existing methods (GxGrare, PLINK-epi, and PLINK-fast). We first inserted the assumed causal variants with a MAF of 1%, a threshold commonly used to define rare variants, and assessed the ability of these four methods to detect causal interactions under three different epistatic models: 1- the threshold model, 2- the multiplicative model, and 3- classic epistasis model. As shown in Figure 2, in all threshold model (Model 1) scenarios, GGI-RUNNER was more powerful than the three alternative methods at the significance level of α = 0.05. For example, GGI-RUNNER achieved a power of 91% when the genotype odds ratio (OR) was 10 and the sample size was 3000 (half cases and half controls). In contrast, the alternative tests had less than 10% power under the same conditions. Increasing the effect size and sample size could further enhance the power of GGI-RUNNER but did not significantly improve the performance of the other three methods. The most powerful alternative method, PLINK-epi, only obtained a power of 38% at the sample size of 4000 with an OR of 20 for causal variants. In the other two epistatic models, the multiplicative model (Model 2) and classic epistatic model (Model 3), GGI-RUNNER still demonstrated the highest power and showed similar increasing trends of power as the sample size and effect size of causal variants increased (Figure 2). The difference in power between GGI-RUNNER and the other three methods was more obvious in Model 1 and Model 2, where the interaction effects are dominant.

**Figure 2.**
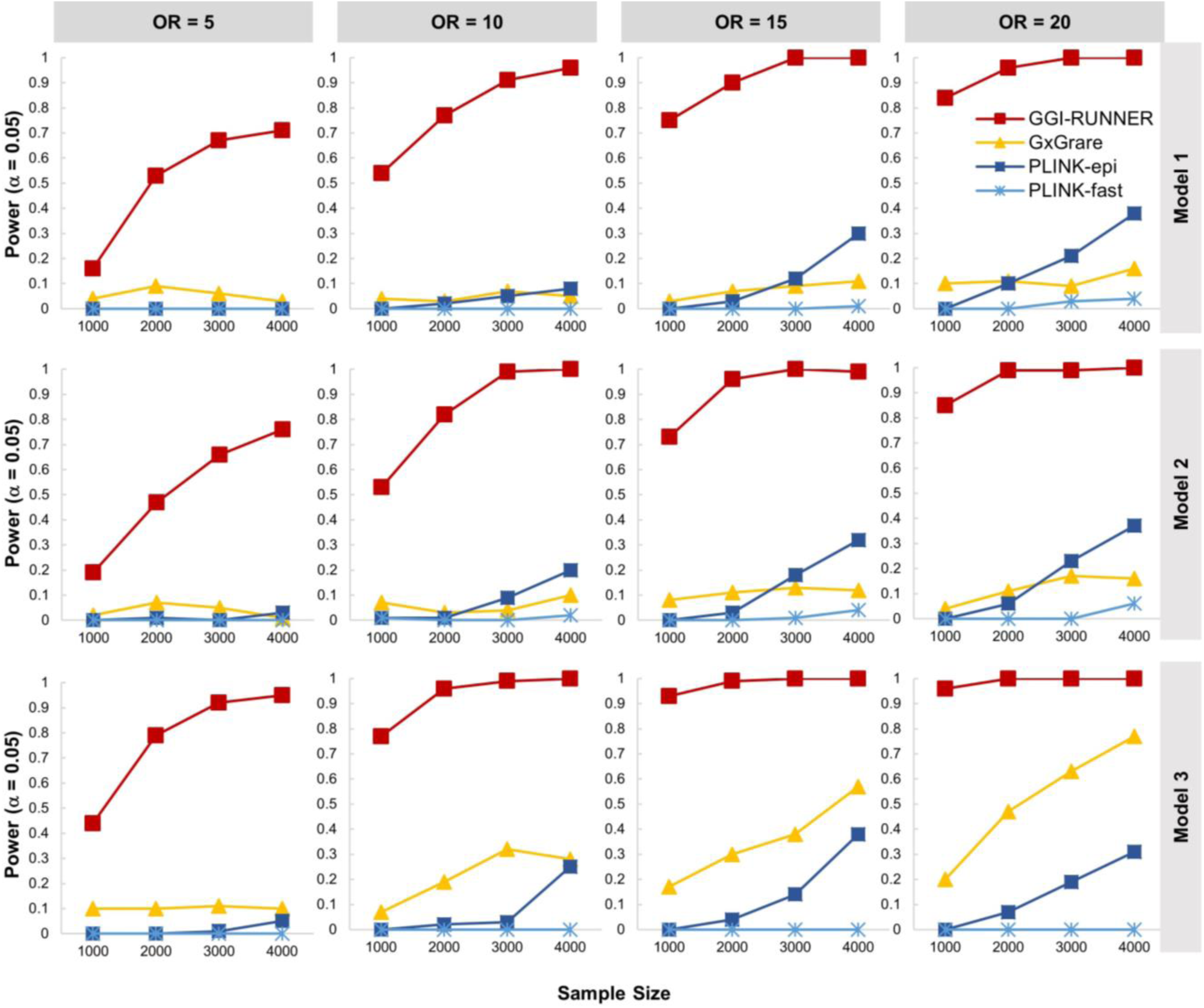
Power curves of four methods: GGI-RUNNER, GxGrare, and two SNP-based epistasis tests implemented in PLINK, ‘--epistasis’ (PLINK-epi) and ‘--fast-epistasis’ (PLINK-fast). Panels from left to right represent scenarios where the genotype odds ratio (OR) for causal variants are 5, 10, 15 and 20, with a fixed disease prevalence of 0.01 and a minor allele frequency (MAF) of 1%. In each plot, the power of the four methods was tested at sample sizes of 1000, 2000, 3000 and 4000 under a balanced case-control design. Power was estimated as the proportion of *P*-values less than 0.05 among 100 replicates. Model 1: the threshold model, Model 2: the multiplicative model, and Model 3: the classic epistasis model.

The power of interaction tests is greatly affected by the number of mutation alleles, which depends on both the sample size and the allele frequency of the variants. To further investigate this, we examined scenarios where causal variants have lower MAFs of 0.1%, 0.25%, 0.5%, or higher MAFs of 2% at a fixed sample size of 2000 cases and 2000 controls. Figure S5 displays the power curves of the four methods, and our method consistently outperformed the other three tests in detecting casual interactions under all three epistatic models. In particular, GGI-RUNNER achieved much higher power than the other methods when the causal variants had a lower MAF of 0.5% at the significance level of α = 0.05. For instance, when the OR was 10 in Model 3, GGI-RUNNER achieved a power of 83%. In comparison, GxGrare only had 9% power, and two pairwise interaction tests implemented in PLINK showed no ability to detect the causal interaction in this scenario. When the MAFs of causal variants were as low as 0.1%, all methods had low power at the tested sample sizes.

However, as the MAFs increased to 0.25%, the power of GGI-RUNNER increased significantly as the OR increased from 5 to 20, particularly in Model 3. In contrast, the other three methods did not recover well in power. When the MAFs of causal variants were as high as 2% in the population, in the simulated case samples, the causal variants could be considered as common variants (MAF > 5%) under the assumed ORs. In this scenario, we observed a significant increase in power for the two PLINK methods. For example, when the OR was 10 in Model 1, PLINK-epi and PLINK-fast achieved the power of 95% and 73%, respectively, but GGI-RUNNER still had the highest power of 100%.

We also explored the power of the four methods at a stricter significance level of α = 1 × 10^−6^, power curves are shown in Figures S6 and S7. Interestingly, using a more stringent threshold did not make a significant difference in power for GGI-RUNNER, which still outperformed the other methods. However, the other three methods nearly lost their power to detect causal interactions at the stricter threshold.

### Application to real datasets of complex diseases

We further applied GGI-RUNNER to identify gene-gene interactions associated with four complex diseases: Alzheimer’s disease (AD), Hirschsprung’s disease (HSCR), Ulcerative colitis (UC) and Crohn’s disease (CD). The first two study samples (AD and HSCR) are all of East Asian ancestry^41; 42^, and the latter two study samples (UC and CD) primarily consist of individuals of European ancestry from the UK Biobank WES data^43^. Rare non-synonymous variants were analyzed in these studies, including missense, start-loss, stop-loss, stop-gain, splicing, frameshift and non-frameshift rare variants with MAFs < 3% in the ancestry-matched panel of gnomAD. Then, GGI-RUNNER was performed on gene pairs with a DIEP score over 0.8 for each of the four complex disease samples.

#### Alzheimer’s disease

Alzheimer’s disease (AD) is a neurodegenerative disorder primarily characterized by cognitive decline and memory loss. The AD dataset consists of 246 Hong Kong Chinese patients with ApoE ɛ4-negative AD and 172 ethnic-matched controls^41^. After quality control at variants and genotypes, a total of 53,769 rare variants in 14,599 genes were retained, and 52,892 pairs of protein-coding genes were included in the interaction analysis (summarized in Table S2). In such a small sample, the overall distribution of GGI-RUNNER *P*-values was still well calibrated as the QQ plot shown in Figure 3A.

**Figure 3.**
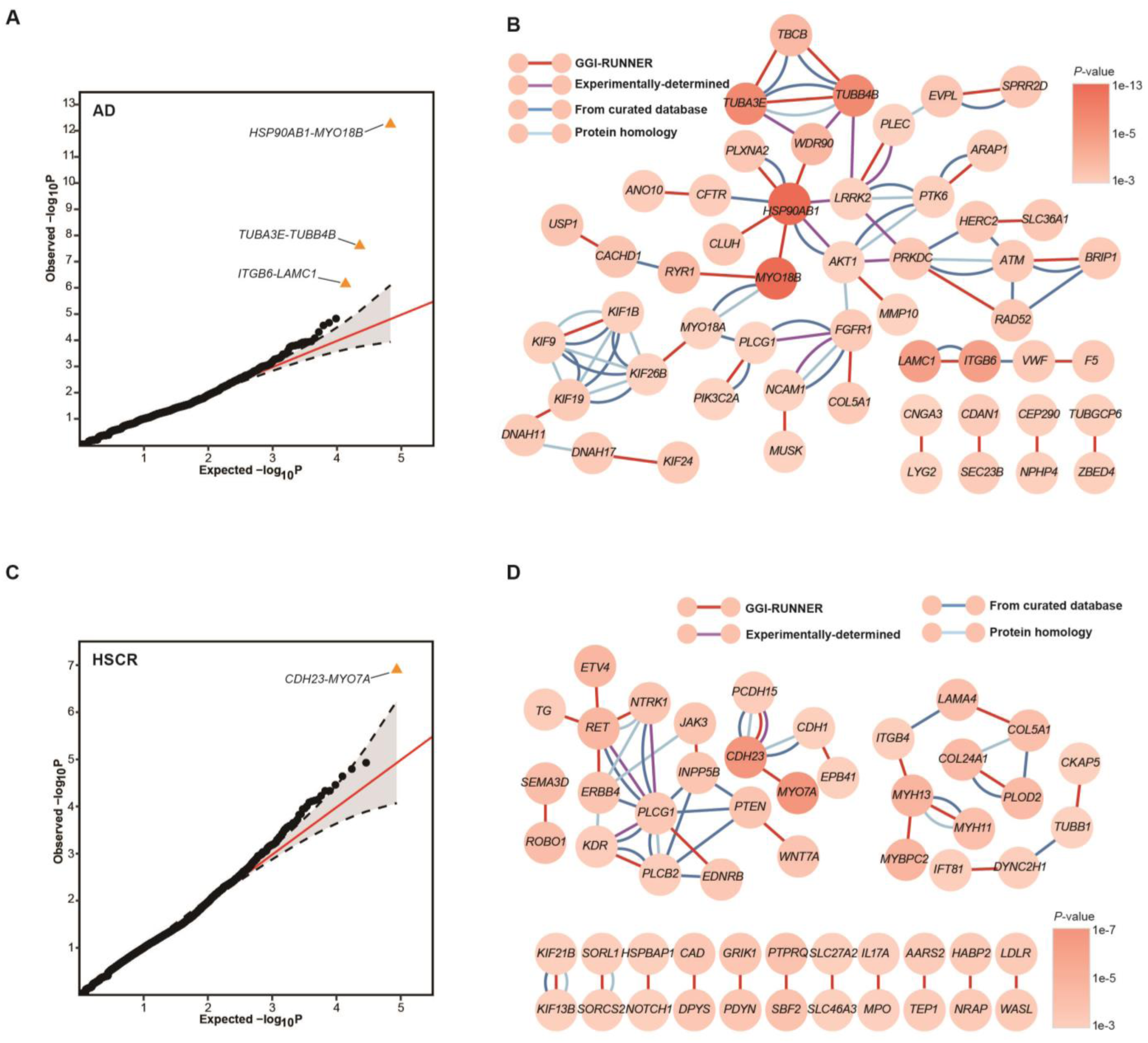
Discovery of gene-gene interactions associated with AD and HSCR by GGI-RUNNER. (A) The QQ plot of *P*-values generated by GGI-RUNNER in AD (246 cases, 172 controls). The orange triangles indicate BHFDR-significant gene-gene interactions (BHFDR *q* <0.05). (B) The network diagram of the top 30 most interactions in GGI-RUNNER analysis for AD. The darker orange nodes correspond to genes with more significant GGI-RUNNER *P*-values. The red edges indicate interactions identified by GGI-RUNNER. Other colored edges originate from interactions with confidence scores ≥0.5 in each category of STRING (purple line for Experimentally determined interactions; dark blue line for interactions from curated databases; light blue line for protein homology), text-mining connections not considered. (C) The QQ plot of *P*-values generated by GGI-RUNNER in HSCR (443 cases, 493 controls). (D) The network diagram of the top 30 most significant interactions in GGI-RUNNER analysis for HSCR.

The ten most significant gene-gene interactions identified by GGI-RUNNER are detailed in Table 3. Remarkably, the most significant result was the interaction between *HSP90AB1* and *MYO18B* genes, yielding a *P*-value of 5.23 × 10^−1^^3^, which also reached study-wide significance (BHFDR *q*-value = 1.77 × 10^−8^). *HSP90AB1*, a member of the Hsp90 family of chaperones, is known to participate in AD pathology by mediating the autophagic clearance of *tau* and *Aβ* aggregates^50^. The second significant gene-gene interaction was *TUBA3E-TUBB4B* (*P* = 2.29 × 10^−8^, *q* = 3.88 × 10^−4^). Both *TUBA3E* and *TUBB4B* are members of the tubulin gene family, encoding for alpha (α)-tubulin and beta (β)-tubulin subunits, respectively. α- and β- tubulin are crucial for microtubule stabilization and have been linked to AD previously^51^. Additionally, we identified another tubulin-related gene, tubulin folding cofactor B (*TBCB*), with suggestive interactions with *TUBA3E* (*P* = 2.02 × 10^−5^) and *TUBB4B* (*P* = 4.56 × 10^−5^). *TBCB* is predicted to play a role in nervous system development and is highly expressed in the brain. The gene pair *ITGB6-LAMC1* also achieved study-wide significance (*P* = 6.50 × 10^−7^, *q* = 7.35 × 10^−3^). *ITGB6* encodes a subunit of the integrin, a family of cell-surface adhesion receptors that is likely to contribute to imbalanced synaptic function in AD^52^. *LAMC1* encodes a member of the laminins, an important extracellular matrix component. Studies have suggested that laminins interact with beta-amyloid peptide (*Aβ*) and contribute to the formation of amyloid plaques, a characteristic feature of AD brains^53^. By integrating known interactions documented in the STRING database, we observed that the top 30 interactions (involving 53 genes) identified by GGI-RUNNER formed a tightly interconnected network (Figure 3B), which holds promising potential for studying the pathways contributing to AD susceptibility.

**Table 3.** Gene-gene interaction analysis on rare variants using GGI-RUNNER and competing methods. Exome-wide gene-gene interaction analyses were performed using GGI-RUNNER on rare non-synonymous variants with a minor allele frequency (MAF) of less than 3% in the East Asian panel of gnomAD. These gene pairs were also tested using three other methods on the same data. The *P*-values of GxGrare were estimated based on 10^6^ permutations. The *P*-values of PLINK-epi (min) and PLINK-fast (min) represent the minimum calculated *P*-values for all possible pairs of rare variants between the two genes using PLINK-epi and PLINK-fast, respectively. /, no *P*-value was obtained in the analysis. Chr. denotes the chromosome number of the human genome where the gene is located.

| Trait | Gene1 | Chr. | Gene2 | Chr. | P-value |  |  |  |
| --- | --- | --- | --- | --- | --- | --- | --- | --- |
|  |  |  |  |  | GGI-RUNNER | GxGrare | PLINK-epi (min) | PLINK-fast (min) |
| AD | *HSP90AB1* | 6 | *MYO18B* | 22 | 5.23E-13 | 0.1040 | 0.989 | 0.1707 |
|  | *TUBA3E* | 2 | *TUBB4B* | 9 | 2.29E-08 | 0.2271 | 0.9884 | 0.2712 |
|  | *ITGB6* | 2 | *LAMC1* | 1 | 6.50E-07 | 0.5598 | 0.996 | 0.3079 |
|  | *HSP90AB1* | 6 | *WDR90* | 16 | 1.40E-05 | 0.0027 | / | 0.2292 |
|  | *TBCB* | 19 | *TUBA3E* | 2 | 2.02E-05 | 0.6577 | / | 0.3479 |
|  | *MYO18B* | 22 | *RYR1* | 19 | 2.57E-05 | 0.3089 | 0.9876 | 0.3077 |
|  | *TBCB* | 19 | *TUBB4B* | 9 | 4.56E-05 | 0.1530 | / | 0.3827 |
|  | *CACHD1* | 1 | *USP1* | 1 | 7.86E-05 | 0.1204 | / | 0.3788 |
|  | *EVPL* | 17 | *SPRR2D* | 1 | 1.05E-04 | 0.0083 | 0.9915 | 0.1023 |
|  | *PRKDC* | 8 | *RAD52* | 12 | 1.09E-04 | 0.1943 | / | 0.2838 |

For the purpose of comparison, we also utilized PLINK-epi, PLINK-fast and GxGrare to analyze the top ten interactions identified by GGI-RUNNER using the same dataset (Table 3). Among the ten gene pairs, GxGrare detected only one interaction, *HSP90AB1*-*WDR90* (*P*_*GxGrare*_ = 2.7 × 10^−3^), that achieved significance at the Bonferroni correction level of 0.05/10 = 5 × 10^−3^. While 8 out of 10 gene pairs did not achieve the classic significant threshold of 0.05 in GxGrare tests. In two SNP-SNP epistasis tests performed by PLINK, PLINK-fast yielded the lowest *P*-value of 0.1023 for the rs7342883-rs143912640 rare variant (RV) pair between *EVPL* and *SPRR2D* genes, and PLINK-epi produced a minimal *P*-value of 0.996 for the gene pair *ITGB6-LAMC1*. The results of PLINK further emphasize the limitations of SNP-SNP epistasis tests in detecting interactions involving rare variants. From another perspective, these findings also suggest that the significant interactions identified by GGI-RUNNER may not be solely driven by individual pairs of RVs but rather by the cumulative effects of multiple RVs interactions between the two genes. Similar results were observed in the other three diseases (Table S4), highlighting the effectiveness of GGI-RUNNER in identifying disease-relevant rare variant interactions that might be missed by other methods.

#### Hirschsprung’s disease

Hirschsprung’s disease (HSCR) is a rare and heterogeneous development disorder of the enteric nervous system (ENS), characterized by the absence of ganglia in the distal colon ^54^. In this study, we analyzed a dataset comprising 443 Asian HSCR patients and 493 controls^42^. The most significant interaction in HSCR was between the *CDH23* and *MYO7A* genes, with a *P*-value of 1.23 × 10^−7^ (BHFDR *q* = 5.35 × 10^−3^, Figure 3C and Table S4). *CDH23* is a member of the cadherin gene family, playing a crucial role in cell-cell adhesion. Abnormalities in neural crest cell development, including those seen in HSCR, often involve alterations in cadherin expression and function^55^. *MYO7A* encodes a myosin protein essential for anchoring cadherins to actin. Previous studies have reported a close interaction between *MYO7A* and *CDH23* in the human Usher syndrome^56^, suggesting their potential interaction in HSCR. Among the top ten interactions identified by GGI-RUNNER (Table S4), we observed three interactions: *EVT4-RET* (*P* = 1.60 × 10^−5^), *ROBO1-SEMA3D* (*P* = 3.48 × 10^−5^), and *ERBB4-RET* (*P* = 7.48 × 10^−5^), involving well-known HSCR risk genes *RET* and *SEMA3D*^54^. When looking at the network formed by the 54 genes involved in the top 30 gene-gene interactions (Figure 3D), we found that these mentioned genes are tightly interconnected within a subnetwork (the left part). Moreover, genes in the subnetwork show highly relevant biological functions to HSCR, including processes such as “neural crest cell development (GO:0055006), differentiation (GO:0014033) and migration (GO:0001755)”, “regulation of nervous system development (GO:0051960)” and “neuromuscular process (GO:0050905)”. Interestingly, we observed that the genes within the right subnetwork primarily belong to the family of cytoskeletal proteins or basement membrane, which appear to implicated in the colonization and/or differentiation of the ENS by regulating cell proliferation, cell-cell adhesion, cell migration and cell projections^57; 58^.

In the discovery phase, we found that all the six genomic features used in the GGI-RUNNER regression model are statistically significant and collectively contributed to the observed approximately uniform distribution of *P*-values (max *P*= 1.54 × 10^−9^) in the HSCR dataset (Table S5). The predictor of RVIB in controls had the lowest significance level (*P*=0.031). In the small AD dataset, this predictor did not even reach statistical significance (*P*=0.399). While in the UC and CD datasets with larger (control) sample sizes, the predictor showed a significant contribution (*P*= 4.30 × 10^−4^ in UC and *P* = 2.57 × 10^−5^ in CD), albeit with small corresponding coefficient values (β =0.082 in UC and β =0.066 in CD). This observation suggests that GGI-RUNNER has little dependence on control samples, especially when the number of control samples is small. This finding also helps explain why the performance of GGI-RUNNER is less affected when there is population stratification among control samples, as mentioned earlier (Table S3 and Figure S4).

#### Two major forms of inflammatory bowel diseases from the UK Biobank WES data

Ulcerative colitis (UC) and Crohn’s disease (CD) are two related yet different forms of inflammatory bowel diseases (IBDs). Here, we utilized UK Biobank WES data^43^ to investigate gene-gene interactions associated with UC and CD using our proposed method. By analyzing rare variants in 1605 cases with CD and 3906 cases with UC from England, along with an equal number of controls, GGI-RUNNER identified three interactions associated with IBDs at a BHFDR of *q* < 5% (Figure 4A, 4B and Table S4). These included *DUOX1-NOD2* and *BMP4-EPHA7* for CD, as well as *ACTRT2-EP400* for UC. Specifically, for CD, GGI-RUNNER identified *DUOX1-NOD2* as the most significant interaction (*P* = 8.71 × 10^−7^). *NOD2* is the first susceptibility gene identified for CD and plays a pivotal role in bacterial infection defense and intestinal epithelium barrier function ^59^. Its interacting gene, *DUOX1*, has also been reported to be involved in innate immune responses to epithelial injury and microbial triggers ^60^. Genes in another notable interaction, *BMP4* and *EPHA7* (*P* = 8.83 × 10^−7^), are implicated in maintaining intestinal barrier integrity and have been associated with chronic inflammatory diseases ^61; 62^. For UC, the most relevant interaction was *ACTRT2-EP400* (*P* = 7.03 × 10^−7^). *ACTRT2* encodes an actin-related protein that mediates cytoskeletal organization, a key process in maintaining intestinal epithelial barrier function ^63^. The E1A binding protein 400 (*EP400*) is a variant chromatin-specific transcription factor that regulates gene activation ^64^. Although there is direct or indirect evidence of these genes participating in intestinal inflammation, as far as we know, their interactions in the context of IBDs have not been previously revealed.

**Figure 4.**
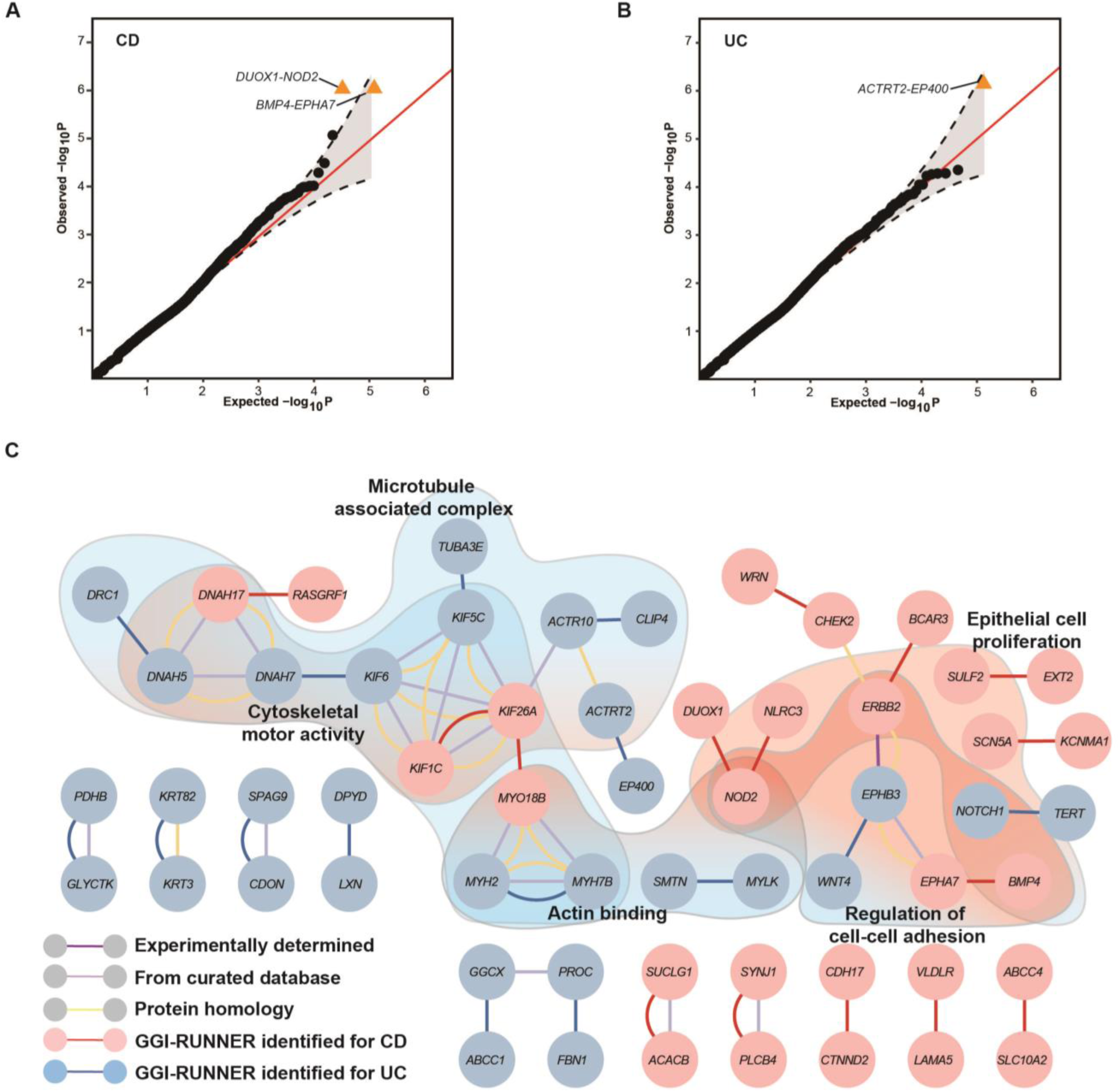
Discovery of gene-gene interactions associated with UC and CD by GGI-RUNNER. (A) The QQ plot of *P*-values generated by GGI-RUNNER in CD (1605 cases, 1605 controls). The orange triangles indicate BHFDR-significant gene-gene interactions (BHFDR *q* <0.05). (B) The QQ plot of *P*-values generated by GGI-RUNNER in UC (3906 cases, 3906 controls). (C) The network diagram of the top 15 significant interactions in GGI-RUNNER analysis for UC and CD. The blue and red nodes correspond to genes identified in UC and CD, respectively. The blue and red edges indicate interactions identified by GGI-RUNNER in UC and CD, respectively. Other colored edges originate from interactions with confidence scores ≥0.5 in each category of STRING (dark purple line for experimentally determined interactions; light purple line for interactions from curated databases; yellow line for protein homology), text-mining connections not considered. Nodes within the shaded areas indicate these genes are enriched in the corresponding GO functions (GO enrichment *q*<0.05).

From the network perspective (Figure 4C), we observed that the top 15 gene-gene interactions identified in UC can be connected to those found in CD through well-documented interactions. In addition, the results of GO enrichment analysis also indicated that the genes prioritized for UC and CD could be enriched in the same GO functions. These findings are consistent with the understanding that susceptibility loci and pathways are shared between UC and CD^65^. However, we also noted that a strikingly large proportion of UC-associated genes enriched in the cytoskeletal-related functions of “cytoskeletal motor activity (GO:0003774)”, “microtubule associated complex (GO:0005875)” and “actin binding (GO:0003779)”. While the CD-associated genes are predominantly enriched in “epithelial cell proliferation (GO:0050673)” and “regulation of cell-cell adhesion (GO:0022407)”. These distinct enrichment patterns suggest that the underlying pathogenesis pathways for UC may differ from those for CD.

### Computation cost

GGI-RUNNER is implemented as a gene-gene interaction analysis module within the KGGSeq software platform, a Java package to identify genetic loci responsible for human diseases or traits by high-throughput sequencing data. The computation time for the GGI-RUNNER depends on the sample size and the number of gene pairs being tested. A whole GGI-RUNNER analysis, including loading and parsing genotypes, quality control, gene feature annotation, and gene-gene interaction tests. On 10 2.40 GHz computing cores with 6-gigabyte memory, GGI-RUNNER took approximately 18 minutes to test interactions for 52,892 gene pairs sequenced on 246 cases and 172 controls. Analyzing 117,012 gene pairs on 7,812 individuals (equal number of cases and controls) required 2.28 hours using the same approach and computational resources (Table S2).

## Discussion

In this study, we introduce a novel statistical framework called GGI-RUNNER, designed to analyze gene-gene interactions involving rare variants in exome-wide association studies of complex diseases. In contrast to the conventional case/control methods, GGI-RUNNER takes a different approach by assessing the enrichment of interactions among rare variants in pairs of genes within patient cohorts. This is achieved by evaluating the extent to which the observed burden of rare variant interactions in cases deviates from the estimated baseline burden in the general population. Through extensive simulations, GGI-RUNNER demonstrated much higher statistical power than alternative methods^17; 18; 66; 67^. Furthermore, it also had its robustness against population stratification. When applied to four representative complex diseases, GGI-RUNNER has revealed, for the first time, a multitude of significant gene-gene interactions contributing to the predisposition of these diseases through rare variants. Importantly, many of these implicated genes would remain undetected using conventional interaction analysis. Notably, the significant gene pairs automatically form intricate and highly interconnected networks, offering unparalleled insights into comprehending the mechanisms underpinning polygenic complex diseases.

The better performance of GGI-RUNNER on interaction analysis of rare variants is first attributed to the adoption of an appropriate estimation model. GGI-RUNNER uses a truncated negative-binomial distribution-based regression model to predict the baseline rare variants interaction burden of pairwise genes which is expected in the general population. A data-driven algorithm allows the model to choose an appropriate truncation point. It is particularly useful to prevent potential jeopardization of model fitting due to the fact that too many gene pairs in the real data have no or low rare variants interaction burden. In the present study, GGI-RUNNER fitting the null model by regression RVIB of background gene pairs on six predictors from publicly available gene/variant annotations and one predictor from the local control samples. We demonstrated that these predictors can lead to an approximately uniform distribution of *P*-values in both simulation and real data analysis. The significance of each predictor changes dynamically from sample to sample (Table S5). The flexible model framework of GGI-RUNNER also allows the usage of other predictors and allows us to study high-order interactions in the near future.

The enhanced power of GGI-RUNNER is also attributed to the flexible, functional weights for rare variants. GGI-RUNNER incorporates integrative functional prediction scores of rare variants to make a more informative RVIB in the regression model. The original functional score is a decimal number ranging from 0 to 1, which needs to be converted into an integer weight to apply to the truncated negative binomial regression model. To set an optimal weighting scale, we conducted a grid search procedure to adjust the scale by minimizing the inflation index of MLFC. While in GxGrare, it arbitrarily uses 0 or 1 as weight values^18^. Furthermore, our method does not need an assumption about the distribution of functional scores. Therefore, it also avoids producing extreme conservatism or optimism testing results resulting from a wrong assumption that does not match the probability distribution of functional scores.

Besides several methodological innovations mentioned above, GGI-RUNNER further enhanced power by reducing the number of tests. The knowledge from external databases, such as STRING^68^, BioGRID^69^, and humanNet^70^, can help us select gene pairs that are more likely to be biologically relevant, enabling more efficient and effective analysis of gene-gene interactions in complex diseases^71^. In our study, GGI-RUNNER used prediction scores from DIEP^34^ by default to narrow all gene pairs down to a subset of pairs with digenic interaction effects potential on disease phenotypes. Users are also allowed to use other external databases to select gene pairs for testing, as long as the data format is a gene-gene-value three-column list. Such a filtering approach can significantly save computational time in genome-wide interaction analysis. The efficient computation of GGI-RUNNER also benefits from the fact that it is a regression-based method. Therefore, it could obtain a *P*-value directly without the need for permutation. This is an attractive feature because it allows for rapid estimation of *P*-values in exome-wide and genome-wide sequencing studies.

Gene-gene interaction analyses of rare variants using GGI-RUNNER identified seven significant associations with four complex traits. These associations included rare variant interactions of *HSP90AB1-MYO18B*, *TUBA3E-TUBB4B*, *ITGB6-LAMC1* and Alzheimer’s disease; *CDH23-MYO7A* interaction and Hirschsprung’s disease; *DUOX1-NOD2* and *BMP4-EPHA7* for Crohn’s disease; and *ACTRT2-EP400* for Ulcerative colitis that were missed by SNP-SNP epistasis tests of PLINK and existing gene-gene interaction test of GxGrare. Early studies have implicated that the function of these genes may contribute to the risk of associated disorders^50–53; 55; 59–63^. Furthermore, the prioritized genes of GGI-RUNNER formed highly interconnected gene-interaction networks for each trait, which may provide a more complete picture to understand and explore the relationship between susceptibility genes and the etiology of complex disease. Intuitively, GGI-RUNNER is expected to discover more significant gene-gene interactions associated with the disease as the sample size increases. However, in practical analysis, the heterogeneity of the disease (e.g., Hirschsprung’s disease), the complexity of population structure (e.g., the UC and CD datasets with multi-ethnic WES data) and other factors will weaken the ability of GGI-RUNNER to detect significant interactions, as well as other methods for interaction analysis.

An important advantage of GGI-RUNNER is that it is less sensitive to population structure. This is primarily due to the unique testing strategy employed by GGI-RUNNER, which essentially calculates the relative difference between the observed RVIB and the estimated baseline RVIB at gene pairs in case samples. Consequently, the influence of population stratification in control samples on the deviation calculation is minimal (Figure S4). The accurate estimation of baseline RVIB by GGI-RUNNER is attributed to the utilization of frequency information from a reference cohort matched to the cases. Therefore, when using frequencies from ancestry-mixed control subjects (Figure S8 and Table S6) or population-unmatched reference cohorts (Figure S9 and Table S7) to model the gene frequency scores, GGI-RUNNER may exhibit a type I error inflation by selecting more significant gene pairs with the population’s private rare variants.

Theoretically, the unique testing strategy allows GGI-RUNNER to perform interaction analysis on case-only samples. However, when control subjects are absent, GGI-RUNNER exhibits an inflation trend in type I error rate as the sample size increases and the MAF threshold becomes more lenient (Figure S3 and Table S8) since we could not use information from the controls to filter out population-private variants. Therefore, GGI-RUNNER should be used with caution for case-only analysis.

Overall, the proposed GGI-RUNNER method is powerful and fast for analyzing gene-gene interactions of rare variants. Its unique statistical strategy of comparing the observed interaction burden to the baseline burden provides an additional approach for testing disease-associated interactions in binary traits. This attribute renders it particularly appealing for studies with limited sample sizes. Remarkably, the networks stemming from the significant gene-gene interactions identified by GGI-RUNNER hold substantial potential for discovering the intricate molecular mechanisms underpinning complex diseases. As public variant annotation databases continue to improve, GGI-RUNNER can be extended for rare variants in noncoding regions, allowing for more comprehensive interaction analysis of complex diseases. GGI-RUNNER is available as a module within our KGGSeq software platform for high-throughput sequencing data analysis, which can be downloaded from our website: http://pmglab.top/kggseq/.

## Data availability

The sequencing data of five panels of the 1000 Genomes Project (1KGP) was downloaded from ftp://ftp.1000genomes.ebi.ac.uk/vol1/ftp/release/20130502/. The UK Biobank analyses were conducted using the UK Biobank resource under applications No.15422 and No.86920. The high-coverage whole genome sequencing data of SG10K is available on https://ega-archive.org with an accession number of EGAS00001003875. The precalculated DIEP scores were downloaded from https://github.com/pmglab/DIEP/tree/main/Dataset/Dataset_S7-CodingDIScores. The functional annotation data are publicly available and were downloaded from the following links: https://gnomad.broadinstitute.org/ (gnomAD); http://database.liulab.science/dbNSFP (dbNSFP). The alternative tools for interaction analysis were downloaded from the following links: http://bibs.snu.ac.kr/software/gxgrare/ (GxGrare); https://www.cog-genomics.org/plink/1.9/ (PLINK, v1.90 beta6.26 Linux64-bit).

## Code availability

GGI-RUNNER is implemented as a module of KGGSeq, available at http://pmglab.top/kggseq/. Source code and additional information to perform simulation and real case analyses are available at GitHub, https://github.com/pmglab/KGGSeq/.

## Acknowledgments

We thank Zhenghan Lian for his insightful discussions and support with data visualization. We appreciate the genomic data provided by Genome Aggregation Database (gnomAD), 1000 Genomes Project, Singapore 10K Genome Project and UK Biobank. This work was supported by the Postdoctoral Science Foundation of China (Grant No. 2022M723660 to HJ); the Basic and Applied Basic Research Foundation of Guangdong Province (Grant No. 2022A1515110913 to HJ); and the National Natural Science Foundation of China (Grant No. 31970650 to ML).

