## Supplemental File for "Exploring Genetic Interactions with Rare Variants Reveals Gene Networks Susceptible to Complex Diseases"

**Supplemental Methods**

**The truncated negative-binomial distribution model**

Given the facts that a large over-dispersion distribution was observed for rare variant interaction burden (RVIB) across protein-coding gene pairs in typical-size samples (Figure S1), we propose to use a truncated negative-binomial (TNB) distribution, where a data-dependent truncation point $t$ is used to limit the range of values for $Y_{ij}$, thus the RVIB statistic $Y_{ij}$ of the gene pair $(i,j)$ as follows:

$$Y_{ij} \sim TNB(\mu_{ij}, \theta, t),\ Y_{ij} = \mathrm{t}+1, \mathrm{t}+2, \cdots,$$

where $\mu_{ij}$ is the expected baseline RVIB of the gene pair $(i,j)$, and $\theta$ is a dispersion parameter. The probability mass function (PMF) of TNB distribution is:

$$g\left(Y_{ij}\middle|\mu_{ij}, \theta, t\right) = \frac{f(Y_{ij}|\mu_{ij}, \theta)}{1-\sum_{s=0}^{t} f(s|\mu_{ij}, \theta)},$$

where $f(Y_{ij}|\mu_{ij}, \theta) = \frac{\Gamma(Y_{ij}+\theta)}{\Gamma(\theta)Y_{ij}!} \frac{\mu_{ij}{}^{Y_{ij}}\theta^{\theta}}{(\mu_{ij}+\theta)^{Y_{ij}+\theta}}$, is the PMF when assuming that $Y_{ij}$ follows a standard negative-binomial distribution $NB(\mu_{ij}, \theta)$, $\Gamma(x)$ is the gamma function, $Y_{ij} = 0,1,2, \ldots$. Similarly, $f(s|\mu_{ij}, \theta)$ is the PMF when $s \sim NB(\mu_{ij}, \theta)$.

**Calculate the deviance residuals and *P*-values**

We use the deviance residuals to measure the deviation of an observed RVIB from the baseline RVIB estimated by the above truncated negative binomial regression. The deviance residual of the model at the gene pair $(i,j)$ is:

$$d_{ij} = sign(e_{ij})\sqrt{2\left|\ln[g(Y_{ij}|\mu_{ij}^{*}, \hat{\theta}, t)] - \ln[g\left(Y_{ij}\middle|\hat{\mu}_{ij}, \hat{\theta}, t\right)]\right|},$$

where $\mathrm{sign}(e_{ij})$ is the sign function of the raw residual $e_{ij}$, $e_{ij} = Y_{ij} - \frac{\hat{\mu}_{ij}}{1-\sum_{s=0}^{t} f(s|\hat{\mu}_{ij}, \hat{\theta})}$, the $\mu_{i}^{*}$ is estimated solution given the observed interaction burden of $Y_i$ and the estimated $\hat{\theta}$ of a saturated model. The $\mu_{ij}^{*}$ can be quickly obtained by solving the equation: $Y_{ij} = \frac{\mu_{ij}^{*}}{1-\sum_{s=0}^{t} f(s|\mu_{ij}^{*}, \hat{\theta})}$.

Then, the deviance residual $d_{ij}$ can be further standardized as $\acute{d}_{ij}$. The corresponding *P*-value for the gene pair $(i,j)$ is approximated by:

$$P_{ij} = 1 - \Phi(\acute{d}_{ij}),$$

where $\Phi(x)$ is the cumulative distribution function of the standard normal distribution.

**Calculate the accumulated rare variants allele frequency for a gene**

Suppose a gene $i$ has $k$ rare variants as defined in a reference population database. Let a rare variant $p$ of the gene $i$ has an allele frequency $f_{i,p}$ and an integer functional weight $w_{i,p}$. Then the accumulated rare variants allele frequency for gene $i$ can be calculated as:

$$X_{2,i} = \sum_{p=1}^{k} f_{i,p} \times w_{i,p}$$

In the present paper, we use the Genome Aggregation Database (gnomAD, V2.1) for variant allele frequency annotation. For a specific studied sample, allele frequencies from the ancestry-matched panel of gnomAD were used for calculation. Variant sites and allele frequencies of seven panels are provided in gnomAD, including East Asian, South Asian, African/African American, Latino, Finnish, Non-Finnish European, and Ashkenazi Jewish. When there are more than one reference population databases, the average allele frequencies from multiple databases are used for the calculation.

**Relationship inference by KING**

We utilized the KING toolset (https://www.kingrelatedness.com/) to assess the sample relationships in the Singapore 10K Genome Project. Following the recommendation of the KING authors, no LD pruning or quality control filtering was performed before conducting the analysis. Using the '—ibdseg' option in KING, we determined the relationship classes for each pair of individuals by analyzing the identical-by-descent (IBD) segments shared between relatives. Individuals with inferred relationships of less than three degrees, including duplicates or monozygotic twins (Dup/MZTwin), parent-offspring pairs (PO), full siblings (FS), and second-degree relatives ($2^{nd}$), were filtered out.

**Two-locus interaction models for power simulation**

We consider three built-in two-locus interaction models whose odds tables are given in Table 1. Model 1 is a threshold model that the odds have a baseline value (α) unless both loci have at least one disease-associated allele, but additional copies of the disease-associated alleles do not increase the risk further. This model can also be called the jointly dominant-dominant model. Model 2 is a multiplicative model that specifies each additional copy of the disease-associated allele at loci A and B further increases the odds in a multiplicative fashion. Model 3 is a classic epistasis model that one allelic effect is blocked by another allele at a different locus. Both loci have the same effect size.

Let $\Pr(\mathrm{D}|g_i)$ denote the probability of an individual being affected given its genotype combination of $g_i$ (i.e., the penetrance of $g_i$), and let $\Pr(\bar{D}|g_i)$ denote the probability of an individual not being affected given its genotype $g_i$. Based on the definition of the odds of disease,

$$ODD_{g_i} = \frac{\Pr(\mathrm{D}|g_i)}{\Pr(\bar{D}|g_i)} = \frac{\Pr(\mathrm{D}|g_i)}{1 - \Pr(\mathrm{D}|g_i)},$$

the penetrance $\Pr(\mathrm{D}|g_i)$ of the genotype $g_i$ can be calculated using the following formula,

$$\Pr(\mathrm{D}|g_i) = \frac{ODD_{g_i}}{1 + ODD_{g_i}}.$$

The disease prevalence $\Pr(D)$ is given as

$$\Pr(D) = \sum_i \Pr(\mathrm{D}|g_i)\Pr(g_i).$$

Once the population prevalence $\Pr(D)$ and the genotype odds ratio $(1 + \theta)$ are fixed in this model, the baseline value α, which indicates the odds of disease when the two loci do not carry any disease alleles, can then be numerically solved based on the above equations.

The frequencies on genotype combinations $\Pr(g_i)$ can be obtained from allele frequencies under the Hardy-Weinberg equilibrium assumption. Once the $3 \times 3$ penetrance table is ready, we can calculate the conditional probability of each genotype given the individual being affected based on the equation:

$$\Pr(g_i|D) = \frac{\Pr\,(g_i)\Pr\,(D|g_i)}{\Pr\,(D)}.$$

### Supplemental Tables

Table S1. The information of assumed risk rare variants in power simulation experiments

**Variant-level**

| Variant | rsID | Gene symbol | MAF1 | MAF2 | FunctionScore |
|---|---|---|---|---|---|
| **1-8025485-G-C** | rs74315353 | *PARK7* | / | 0.0006418 | 0.8468 |
| **1-20966422-G-T** | rs746882245 | *PINK1* | 0.000164 | / | 0.9513 |

**Gene-level**

| Gene | Chr. no. | Start Position | End Position | CDS_Length | OE_mis | OE_lof | ExonGC | DIEP_score |
|---|---|---|---|---|---|---|---|---|
| ***PARK7*** | 1 | 8014351 | 8045565 | 0.567 | 0.9556 | 0.1115 | 0.4951 | 0.9881 |
| ***PINK1*** | 1 | 20959948 | 20978004 | 1.786 | 0.9795 | 0.7659 | 0.5543 | |

The physical positions of variants and genes are given in the UCSC hg19 version of the human genome. Abbreviation: MAF1, the minor allele frequency from the East Asian panel of gnomAD exomes; MAF2, the minor allele frequency from the East Asian panel of gnomAD genomes. Chr. no., chromosome number.

Table S2. Real high-throughput sequencing datasets used in the present paper

| Trait | Description | Number of patients | Number of controls | Source | Human genome version | Initial Variants after QC | Rare non-synonymous variants [a] | Number of tested genes | Number of tested gene pairs [b] | Computation time (min.) [c] |
|---|---|---|---|---|---|---|---|---|---|---|
| **AD** | ApoE ε4-negative Alzheimer's disease | 246 | 172 | Hong Kong. China | Hg19 | 487252 | 53769 | 14599 | 52892 | 18.02 |
| **HSCR** | Hirschsprung's disease | 443 | 493 | Asian | Hg19 | 336307 | 96278 | 16522 | 70895 | 21.43 |
| **CD** | Crohn's disease (ICD10 code: K50) from UK biobank WES data | 1605 | 1605 | England | Hg38 | 2447229 | 380144 | 17775 | 100825 | 79.07 |
| **UC** | Ulcerative colitis (ICD10 code: K51) from UK biobank WES data | 3906 | 3906 | England | Hg38 | 3474384 | 484199 | 17812 | 117012 | 136.95 |

[a] Rare non-synonymous variants, including missense, start-loss, stop-loss, stop-gain, splicing, frameshift and non-frameshift variants, with minor allele frequency<3% in reference populations (East Asian panel of the gnomAD for AD and HSCR, and Non-Finnish European panel of the gnomAD for CD and UC) were analyzed for each dataset. [b] Protein-coding gene pairs with a DIEP score over 0.8 were tested for gene-gene interaction. [c] Computation time in each cell refers to the runtime for the whole procedure, including loading and parsing genotypes, quality control (QC), gene feature annotation, and gene-gene interaction tests.

Table S3. Empirical type I error rates for GGI-RUNNER in ancestral mixed samples.

| Group | Sample size | alpha | | | Number of tested gene pairs | MLFC |
|---|---|---|---|---|---|---|
| | | 1.0E-02 | 1.0E-03 | 1.0E-04 | | |
| **AFR-AMR** | 881 | 6.691E-03 | 6.633E-04 | 8.652E-05 | 34673 | 0.070 |
| **AFR-EAS** | 881 | 6.825E-03 | 5.819E-04 | 7.936E-05 | 37804 | 0.063 |
| **AFR-EUR** | 881 | 6.820E-03 | 6.275E-04 | 8.184E-05 | 36656 | 0.068 |
| **AFR-SAS** | 881 | 6.890E-03 | 6.485E-04 | 5.404E-05 | 37008 | 0.062 |
| **AMR-AFR** | 462 | 5.486E-03 | 8.572E-04 | 2.286E-04 | 17499 | 0.098 |
| **AMR-EAS** | 462 | 6.102E-03 | 8.177E-04 | 1.573E-04 | 31795 | 0.072 |
| **AMR-EUR** | 462 | 5.739E-03 | 7.127E-04 | 7.502E-05 | 26661 | 0.086 |
| **AMR-SAS** | 462 | 6.309E-03 | 9.889E-04 | 3.069E-04 | 29325 | 0.094 |
| **EAS-AFR** | 672 | 6.588E-03 | 9.087E-04 | 4.543E-05 | 22010 | 0.112 |
| **EAS-AMR** | 672 | 6.007E-03 | 7.855E-04 | 4.621E-05 | 21641 | 0.105 |
| **EAS-EUR** | 672 | 8.065E-03 | 1.367E-03 | 1.367E-04 | 21948 | 0.117 |
| **EAS-SAS** | 672 | 7.313E-03 | 8.850E-04 | 4.658E-05 | 21468 | 0.105 |
| **EUR-AFR** | 670 | 5.917E-03 | 6.801E-04 | 2.040E-04 | 14703 | 0.123 |
| **EUR-AMR** | 670 | 4.812E-03 | 7.467E-04 | 8.297E-05 | 12053 | 0.159 |
| **EUR-EAS** | 670 | 5.704E-03 | 5.704E-04 | 1.901E-04 | 15779 | 0.125 |
| **EUR-SAS** | 670 | 6.413E-03 | 4.008E-04 | 6.680E-05 | 14970 | 0.127 |
| **SAS-AFR** | 652 | 5.817E-03 | 6.375E-04 | 3.984E-05 | 25098 | 0.075 |
| **SAS-AMR** | 652 | 6.178E-03 | 6.348E-04 | 0 | 23631 | 0.074 |
| **SAS-EAS** | 652 | 6.318E-03 | 7.152E-04 | 0 | 25167 | 0.102 |
| **SAS-EUR** | 652 | 6.256E-03 | 7.138E-04 | 0 | 23817 | 0.097 |

The 20 samples were simulated based on five population panels of the 1KGP. The five panels are AFR- African, AMR- American, EAS- East Asian, EUR- European, and SAS- South Asian. AFR-AMR in the table denotes that the pseudo-cases in this sample were from the AFR panel, while half of the controls were from the AFR panel, and the other half were from the AMR panel. Same as the other 19 group labels. In these analyses, the MAFs from the reference panel of gnomAD matched to the racial background of the cases (e.g., African/African American (AFR) panel of gnomAD for AFR-AMR) are used for rare variant selection and the accumulated MAF calculation. MLFC: the mean log fold change.

Table S4. Gene-gene interaction analysis on rare variants using GGI-RUNNER and competing methods

| Trait | Gene1 | Chr. | Gene2 | Chr. | *P*-value | | | |
|---|---|---|---|---|---|---|---|---|
| | | | | | GGI-RUNNER | GxGrare | PLINK-epi (min) | PLINK-fast (min) |
| **HSCR** | *CDH23* | 10 | *MYO7A* | 11 | 1.23E-07 | 0.8081 | 0.9912 | 0.2559 |
| | *MYBPC2* | 19 | *MYH13* | 17 | 1.18E-05 | 0.8885 | / | 0.2017 |
| | *ETV4* | 17 | *RET* | 10 | 1.60E-05 | 0.0134 | 0.9913 | 0.0988 |
| | *COL24A1* | 1 | *PLOD2* | 3 | 2.28E-05 | 0.1521 | / | 0.3091 |
| | *ROBO1* | 3 | *SEMA3D* | 7 | 3.48E-05 | 0.9180 | / | 0.4737 |
| | *COL5A1* | 9 | *LAMA4* | 6 | 4.64E-05 | 0.6630 | 0.9964 | 0.3468 |
| | *MYH11* | 16 | *MYH13* | 17 | 4.65E-05 | 0.8354 | 0.9974 | 0.2790 |
| | *PTPRQ* | 12 | *SBF2* | 11 | 5.92E-05 | 1 | / | 0.1812 |
| | *PTEN* | 10 | *WNT7A* | 3 | 7.15E-05 | 0.6126 | / | 0.5617 |
| | *ERBB4* | 2 | *RET* | 10 | 7.48E-05 | 0.0066 | / | 0.2010 |
| **UC** | *ACTRT2* | 1 | *EP400* | 12 | 7.03E-07 | 0.949 | 0.4855 | 0.2246 |
| | *DNAH5* | 5 | *DRC1* | 2 | 4.41E-05 | 0.179 | 0.9889 | 0.0532 |
| | *EPHB3* | 3 | *WNT4* | 1 | 5.20E-05 | 0.268 | 0.996 | 0.3448 |
| | *KIF5C* | 2 | *TUBA3E* | 2 | 5.24E-05 | 0.039 | / | 0.3329 |
| | *DPYD* | 1 | *LXN* | 3 | 5.45E-05 | 0.371 | 0.3541 | 0.2143 |
| | *MYLK* | 3 | *SMTN* | 22 | 5.87E-05 | 0.112 | 0.362 | 0.1669 |
| | *ACTR10* | 14 | *CLIP4* | 2 | 8.88E-05 | 0.648 | / | 0.2246 |
| | *MYH2* | 17 | *MYH7B* | 20 | 8.94E-05 | 0.734 | 0.1808 | 0.1593 |
| | *CDON* | 11 | *SPAG9* | 17 | 1.18E-04 | 0.632 | 0.4470 | 0.1847 |
| | *FBN1* | 15 | *PROC* | 2 | 1.39E-04 | 0.170 | / | 0.1896 |
| **CD** | *DUOX1* | 15 | *NOD2* | 16 | 8.71E-07 | 0.406 | 0.9960 | 0.1650 |
| | *BMP4* | 14 | *EPHA7* | 6 | 8.83E-07 | 0.441 | / | 0.3643 |
| | *PLCB4* | 20 | *SYNJ1* | 21 | 8.55E-06 | 0.889 | / | 0.2130 |
| | *BCAR3* | 1 | *ERBB2* | 17 | 3.26E-05 | 0.277 | 0.3979 | 0.2519 |
| | *KIF26A* | 14 | *MYO18B* | 22 | 5.16E-05 | 0.702 | 0.3088 | 0.2851 |
| | *KCNMA1* | 10 | *SCN5A* | 3 | 9.66E-05 | 0.729 | / | 0.3262 |
| | *KIF1C* | 17 | *KIF26A* | 14 | 9.72E-05 | 0.823 | 0.4348 | 0.1937 |
| | *DNAH17* | 17 | *RASGRF1* | 15 | 1.02E-04 | 0.729 | 0.9879 | 0.1468 |
| | *ABCC4* | 13 | *SLC10A2* | 13 | 1.03E-04 | 0.012 | 0.1811 | 0.2373 |
| | *EXT2* | 11 | *SULF2* | 20 | 1.11E-04 | 0.526 | 0.9894 | 0.2908 |

Exome-wide gene-gene interaction analyses were performed using GGI-RUNNER on rare non-synonymous variants with a minor allele frequency (MAF) of less than 3% in the ancestry-matched panel of gnomAD. The top ten significant interactions identified by GGI-RUNNER for each trait are listed in the table. These gene pairs were also tested using three other methods on the same data. The *P*-values of GxGrare were estimated based on $10^6$ permutations for HSCR, and $10^3$ permutations for UC and CD (In these two traits with larger sample sizes, it took several hours (>7h) to calculate a *P*-value of GxGrare based on $10^6$ permutations for each pair of genes). The *P*-values of PLINK-epi (min) and PLINK-fast (min) represent the minimum calculated *P*-values for all possible pairs of rare variants between the two genes using PLINK-epi and PLINK-fast, respectively. /, no *P*-value was obtained in the analysis. Chr. denotes the chromosome number of the human genome where the gene is located.

Table S5. The estimated parameters of seven predictors by GGI-RUNNER.

| Trait | Variable | Estimate | Std. Error | z value | Pr(>\|z\|) |
|---|---|---|---|---|---|
| **AD** | (Intercept) | 0.9106 | 2.66E-02 | 34.2490 | 4.51E-257 |
| | RegionLength | 0.0045 | 2.28E-04 | 19.8398 | 1.35E-87 |
| | EAS | 9.2274 | 3.76E-01 | 24.5332 | 6.53E-133 |
| | RegionLength×EAS | -0.0258 | 3.05E-03 | -8.4615 | 2.64E-17 |
| | oe_mis | 0.0144 | 2.37E-02 | 0.6106 | 0.5415 |
| | oe_lof | 0.1324 | 2.30E-02 | 5.7577 | 8.52E-09 |
| | ExonGC | 0.3720 | 7.66E-02 | 4.8540 | 1.21E-06 |
| | CtrlRVIB | 0.0526 | 6.23E-02 | 0.8442 | 0.3986 |
| **HSCR** | (Intercept) | 1.1340 | 2.15E-02 | 52.6435 | 0 |
| | RegionLength | 0.0065 | 1.89E-04 | 34.2458 | 5.04E-257 |
| | EAS | 3.3332 | 1.46E-01 | 22.8122 | 3.47E-115 |
| | RegionLength×EAS | -0.0150 | 1.35E-03 | -11.1033 | 1.21E-28 |
| | oe_mis | 0.1620 | 1.93E-02 | 8.4060 | 4.24E-17 |
| | oe_lof | 0.1101 | 1.82E-02 | 6.0396 | 1.54E-09 |
| | ExonGC | 0.7113 | 6.11E-02 | 11.6494 | 2.31E-31 |
| | CtrlRVIB | 0.0581 | 2.69E-02 | 2.1565 | 0.0310 |
| **UC** | (Intercept) | 1.1471 | 1.70E-02 | 67.4594 | 0 |
| | RegionLength | 0.0088 | 1.83E-04 | 47.9856 | 0 |
| | NFE | 6.7303 | 2.04E-01 | 33.0467 | 1.74E-239 |
| | RegionLength×NFE | -0.0295 | 1.86E-03 | -15.9064 | 5.72E-57 |
| | oe_mis | 0.1604 | 1.53E-02 | 10.5129 | 7.53E-26 |
| | oe_lof | 0.0538 | 1.57E-02 | 3.4172 | 6.33E-04 |
| | ExonGC | 0.7429 | 4.89E-02 | 15.1798 | 4.81E-52 |
| | CtrlRVIB | 0.0821 | 2.33E-02 | 3.5213 | 4.30E-04 |
| **CD** | (Intercept) | 1.1417 | 1.97E-02 | 57.9624 | 0 |
| | RegionLength | 0.0070 | 2.02E-04 | 34.7331 | 2.50E-264 |
| | NFE | 8.6024 | 2.57E-01 | 33.4217 | 6.62E-245 |
| | RegionLength×NFE | -0.0314 | 2.18E-03 | -14.3967 | 5.43E-47 |
| | oe_mis | 0.0773 | 1.77E-02 | 4.3704 | 1.24E-05 |
| | oe_lof | 0.0806 | 1.80E-02 | 4.4812 | 7.42E-06 |
| | ExonGC | 0.7645 | 5.63E-02 | 13.5713 | 5.92E-42 |
| | CtrlRVIB | 0.0656 | 1.56E-02 | 4.2087 | 2.57E-05 |

RegionLength: $X_1$, the coding regions (CDS) length. EAS (or NFE): $X_2$, the accumulated MAF calculated based on reference MAFs from the East Asian panel of gnomAD (or the Non-Finnish European panel of gnomAD). RegionLength×EAS (or RegionLength×NFE): $X_3$, the product of CDS length and the accumulated MAF. oe_mis and oe_lof: the observed/expected ratios for missense (oe_mis, $X_4$)and loss-of-function variations (oe_lof, $X_5$). ExonGC: $X_6$ the GC content in exon regions of the gene. $X_1$-$X_6$ are calculated as the product of the feature values of two genes for the gene-gene interaction analysis of GGI-RUNNER. CtrlRVIB: the rare variant interaction burden of gene pairs calculated in the control samples.

Table S6. Empirical type I error rates for GGI-RUNNER using MAF from ancestral mixed controls.

| Group | Sample size | alpha | | | Number of tested gene pairs | MLFC |
|---|---|---|---|---|---|---|
| | | 1.0E-02 | 1.0E-03 | 1.0E-04 | | |
| **AFR-AMR** | **881** | 6.720E-03 | 6.922E-04 | 1.442E-04 | 34673 | 0.076 |
| **AFR-EAS** | **881** | 6.454E-03 | 6.349E-04 | 1.058E-04 | 37804 | 0.074 |
| **AFR-EUR** | **881** | 6.629E-03 | 6.547E-04 | 1.091E-04 | 36656 | 0.075 |
| **AFR-SAS** | **881** | 6.539E-03 | 5.945E-04 | 1.081E-04 | 37008 | 0.065 |
| **AMR-AFR** | **462** | 6.172E-03 | 1.086E-03 | 4.000E-04 | 17499 | 0.129 |
| **AMR-EAS** | **462** | 6.794E-03 | 9.750E-04 | 1.258E-04 | 31795 | 0.077 |
| **AMR-EUR** | **462** | 5.851E-03 | 1.050E-03 | 1.125E-04 | 26661 | 0.106 |
| **AMR-SAS** | **462** | 7.025E-03 | 1.194E-03 | 2.046E-04 | 29325 | 0.094 |
| **EAS-AFR** | **672** | 7.042E-03 | 9.087E-04 | 4.543E-05 | 22010 | 0.113 |
| **EAS-AMR** | **672** | 6.700E-03 | 6.469E-04 | 4.621E-05 | 21641 | 0.117 |
| **EAS-EUR** | **672** | 7.882E-03 | 8.657E-04 | 4.556E-05 | 21948 | 0.100 |
| **EAS-SAS** | **672** | 7.639E-03 | 9.782E-04 | 1.397E-04 | 21468 | 0.100 |
| **EUR-AFR** | **670** | 6.257E-03 | 8.842E-04 | 2.040E-04 | 14703 | 0.156 |
| **EUR-AMR** | **670** | 6.720E-03 | 1.327E-03 | 3.319E-04 | 12053 | 0.185 |
| **EUR-EAS** | **670** | 5.704E-03 | 7.605E-04 | 2.535E-04 | 15779 | 0.167 |
| **EUR-SAS** | **670** | 7.081E-03 | 9.352E-04 | 1.336E-04 | 14970 | 0.136 |
| **SAS-AFR** | **652** | 6.096E-03 | 5.977E-04 | 0 | 25098 | 0.101 |
| **SAS-AMR** | **652** | 6.432E-03 | 7.617E-04 | 4.232E-05 | 23631 | 0.111 |
| **SAS-EAS** | **652** | 7.152E-03 | 9.139E-04 | 7.947E-05 | 25167 | 0.113 |
| **SAS-EUR** | **652** | 7.600E-03 | 7.558E-04 | 4.199E-05 | 23817 | 0.117 |

The 20 samples were simulated based on five population panels of the 1KGP. The five panels are AFR- African, AMR- American, EAS- East Asian, EUR- European, and SAS- South Asian. AFR-AMR in the table denotes that the pseudo-cases in this sample were from the AFR panel, while half of the controls were from the AFR panel, and the other half were from the AMR panel. Same as the other 19 group labels. In these analyses, the MAFs from the reference panel of gnomAD matched to the racial background of the cases (e.g., African/African American (AFR) panel of gnomAD for AFR-AMR) are used for rare variant selection. The MAFs from the ancestral mixed controls are used for the accumulated MAF calculation. MLFC: the mean log fold change.

Table S7. Empirical type I error rates for GGI-RUNNER using MAF from ancestry-unmatched reference.

| Group | Sample size | alpha | | | Number of tested gene pairs | MLFC |
|---|---|---|---|---|---|---|
| | | 1.0E-02 | 1.0E-03 | 1.0E-04 | | |
| AFR-AMR | 881 | 7.561E-03 | 8.335E-04 | 1.389E-04 | 50390 | 0.062 |
| AFR-EAS | 881 | 9.304E-03 | 1.127E-03 | 2.902E-04 | 58577 | 0.055 |
| AFR-EUR | 881 | 8.058E-03 | 1.058E-03 | 2.997E-04 | 56715 | 0.049 |
| AFR-SAS | 881 | 7.740E-03 | 8.580E-04 | 1.926E-04 | 57107 | 0.045 |
| AMR-AFR | 462 | 7.424E-03 | 1.061E-03 | 4.419E-05 | 22629 | 0.072 |
| AMR-EAS | 462 | 8.624E-03 | 1.277E-03 | 2.688E-04 | 44645 | 0.081 |
| AMR-EUR | 462 | 5.952E-03 | 5.653E-04 | 6.651E-05 | 30072 | 0.082 |
| AMR-SAS | 462 | 7.656E-03 | 1.074E-03 | 1.377E-04 | 36313 | 0.072 |
| EAS-AFR | 672 | 7.235E-03 | 8.832E-04 | 1.019E-04 | 29440 | 0.076 |
| EAS-AMR | 672 | 6.756E-03 | 6.965E-04 | 1.741E-04 | 28715 | 0.071 |
| EAS-EUR | 672 | 8.009E-03 | 9.163E-04 | 1.697E-04 | 29466 | 0.083 |
| EAS-SAS | 672 | 8.377E-03 | 7.776E-04 | 1.767E-04 | 28292 | 0.086 |
| EUR-AFR | 670 | 7.057E-03 | 9.703E-04 | 8.821E-05 | 22674 | 0.109 |
| EUR-AMR | 670 | 5.113E-03 | 7.601E-04 | 1.382E-04 | 14472 | 0.147 |
| EUR-EAS | 670 | 6.885E-03 | 1.181E-03 | 2.444E-04 | 24546 | 0.126 |
| EUR-SAS | 670 | 7.277E-03 | 7.814E-04 | 1.954E-04 | 20475 | 0.092 |
| SAS-AFR | 652 | 6.271E-03 | 7.754E-04 | 1.011E-04 | 29661 | 0.117 |
| SAS-AMR | 652 | 6.821E-03 | 7.785E-04 | 1.483E-04 | 26975 | 0.089 |
| SAS-EAS | 652 | 6.619E-03 | 7.426E-04 | 6.457E-05 | 30973 | 0.087 |
| SAS-EUR | 652 | 6.946E-03 | 7.311E-04 | 1.462E-04 | 27355 | 0.110 |

The 20 samples were simulated based on five population panels of the 1KGP. The five panels are AFR- African, AMR- American, EAS- East Asian, EUR- European, and SAS- South Asian. AFR-AMR in the table denotes that the pseudo-cases in this sample were from the AFR panel, while half of the controls were from the AFR panel, and the other half were from the AMR panel. Same as the other 19 group labels. In these analyses, the MAFs from the reference panel of gnomAD matched to the racial background of the half ancestral mixed controls (e.g., Latino (AMR) panel of gnomAD for AFR-AMR) are used for rare variant selection and the accumulated MAF calculation. MLFC, the mean log fold change.

Table S8. Empirical type I error rates for GGI-RUNNER in case-only samples.

| Sample size | MAF cutoff | alpha | | | Number of tested gene pairs | MLFC |
|---|---|---|---|---|---|---|
| | | 1.0E-03 | 1.0E-04 | 1.0E-05 | | |
| 500 | 1% | 6.131E-04 | 9.783E-05 | 1.926E-05 | 55048±817 | 0.085±0.011 |
| 1000 | 1% | 1.095E-03 | 2.831E-04 | 9.979E-05 | 82530±683 | 0.094±0.010 |
| 1500 | 1% | 1.558E-03 | 4.784E-04 | 2.004E-04 | 101246±524 | 0.126±0.012 |
| 2000 | 1% | 1.856E-03 | 6.161E-04 | 2.780E-04 | 115365±408 | 0.168±0.013 |
| 2000 | 0.1% | 5.284E-04 | 6.201E-05 | 5.282E-06 | 56927±371 | 0.069±0.011 |
| 2000 | 0.5% | 1.440E-03 | 4.625E-04 | 1.948E-04 | 100960±460 | 0.126±0.011 |
| 2000 | 5% | 5.158E-03 | 2.186E-03 | 1.096E-03 | 144347±352 | 0.651±0.014 |

GGI-RUNNER was applied to samples with different sample sizes of cases, 500, 1000, 1500 and 2000, and tested gene-gene interactions involving rare variants under different MAF cutoffs, 0.1%, 0.5% and 5%. The semi-simulation procedure produced the samples based on whole-genome sequencing data from SG10K unrelated subjects, the same samples used in Table 1, but not with control subjects. Each setting repeats 100 times. MLFC: the mean log fold change.

**Supplemental Figures**

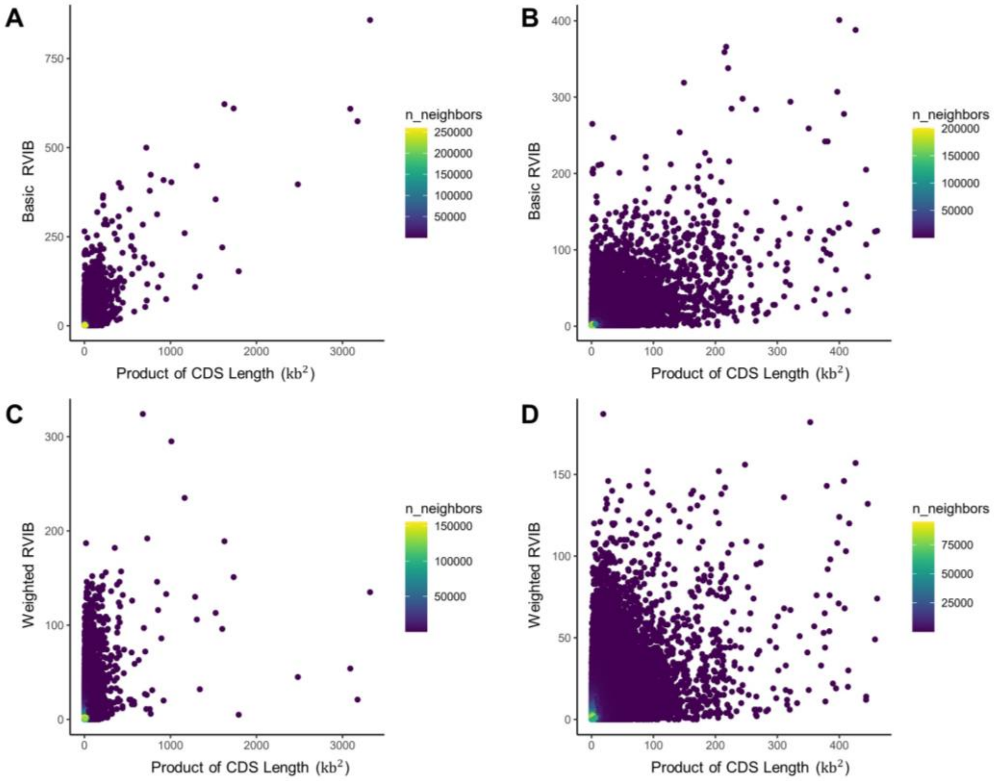


Figure S1. The density scatter plots of the rare variant interaction burden (RVIB) and the gene pair's product of coding regions length (kb$^2$). (A) basic RVIB of all available gene pairs (301457 pairs); (B) basic RVIB of gene pairs with a product of coding region length less than 500 (301419 pairs); (C) functional weighted RVIB of all available gene pairs; and (D) functional weighted RVIB of gene pairs with a product of coding region length less than 500. The RVIB scores were calculated based on 2000 pseudo-cases randomly drawn from the SG10K unrelated subjects. Gene pairs with a DIEP score over 0.8 and an RVIB score over zero were shown in the plots. Each point in the plot is colored by the number of neighboring points.

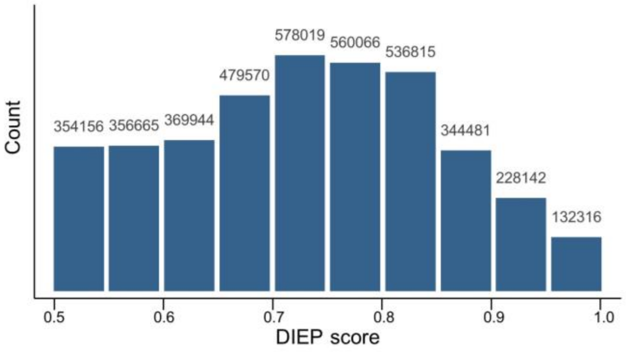


Figure S2. The histogram of protein-coding gene pairs with a DIEP score over 0.5.

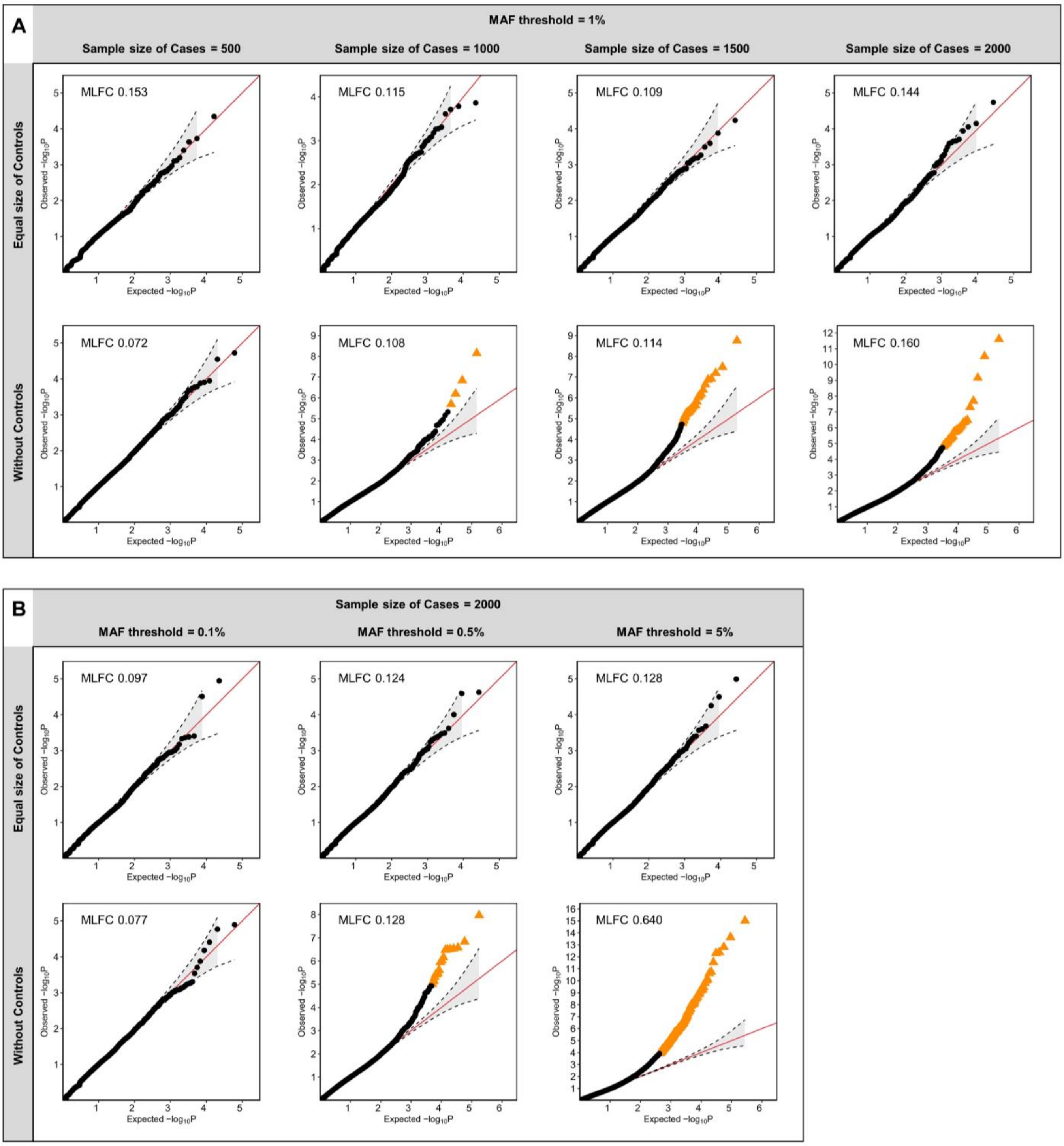


Figure S3. The QQ plots of *P*-values generated by GGI-RUNNER in balanced case-control and case-only samples. (A) GGI-RUNNER was applied to samples with different sample sizes of cases, 500, 1000, 1500 and 2000, along with an equal number of controls or without controls, respectively. The MAF threshold in these analyses was set as 1%. (B) GGI-RUNNER was tested using different MAF cutoffs of 0.1%, 0.5% and 5% in samples containing 2000 cases, along with an equal number of controls or without controls, respectively. The semi-simulation procedure produced the samples based on whole-genome sequencing data from SG10K unrelated subjects. The QQ plots were generated based on the results randomly selected once from 100 simulations in each group. MLFC: the mean log fold change.

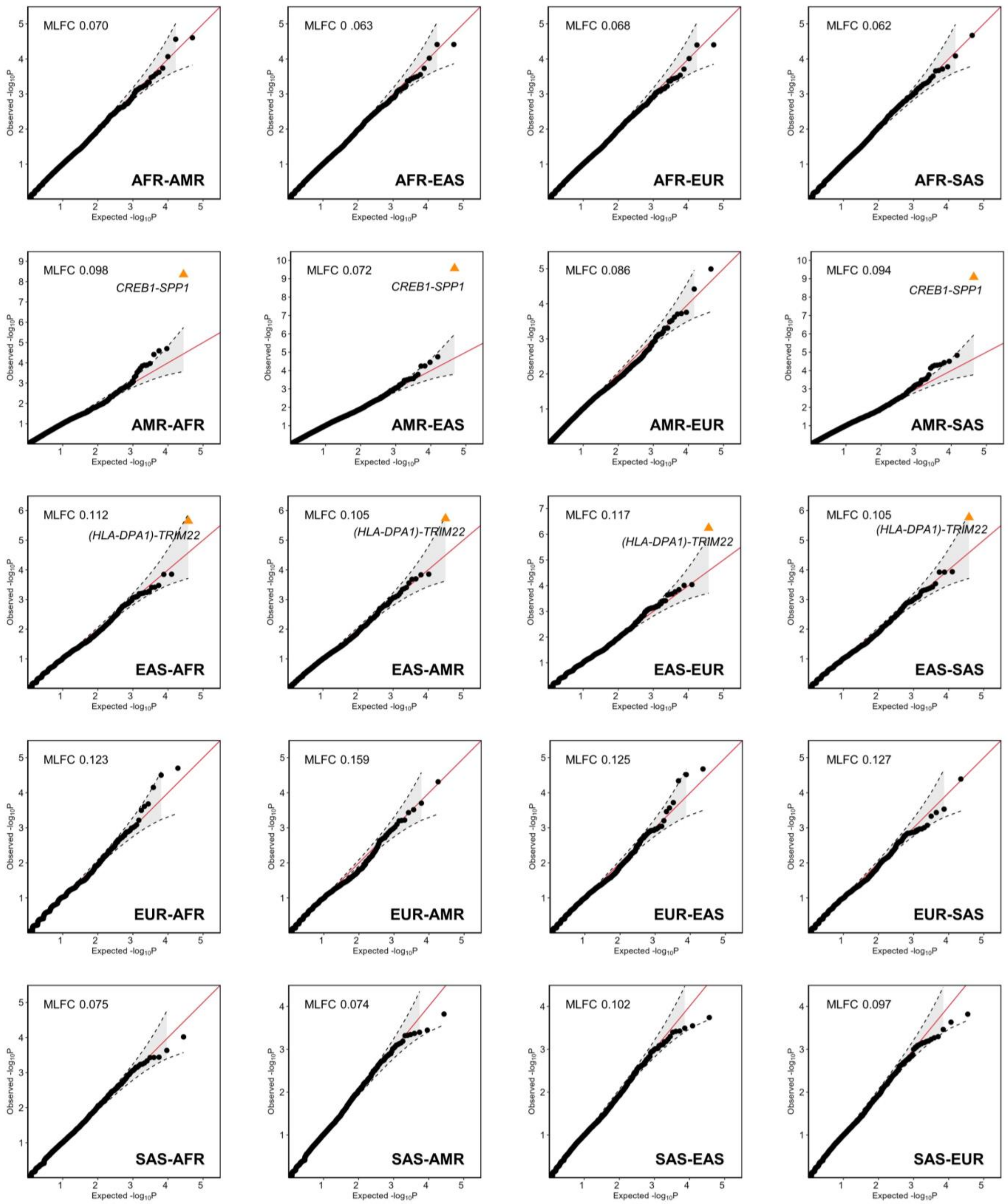


Figure S4. The QQ plots of *P*-values generated by GGI-RUNNER in ancestral mixed samples. The 20 samples were simulated based on five population panels of the 1KGP. The five panels are AFR- African, AMR- American, EAS- East Asian, EUR- European, and SAS- South Asian. AFR-AMR in the table denotes that the pseudo-cases in this sample were from the AFR panel, while half of the controls were from the AFR panel, and the other half were from the AMR panel. Same as the other 19 group labels. In these analyses, the MAFs from the reference panel of gnomAD matched to the racial background of the cases (e.g., African/African American (AFR) panel of gnomAD for AFR-AMR) are used for rare variant selection and the accumulated MAF calculation. The gene symbols of gene pair reaching exome-wide significance (BHFDR $q<0.05$) were annotated below the corresponding point in the plots. MLFC: the mean log fold change.

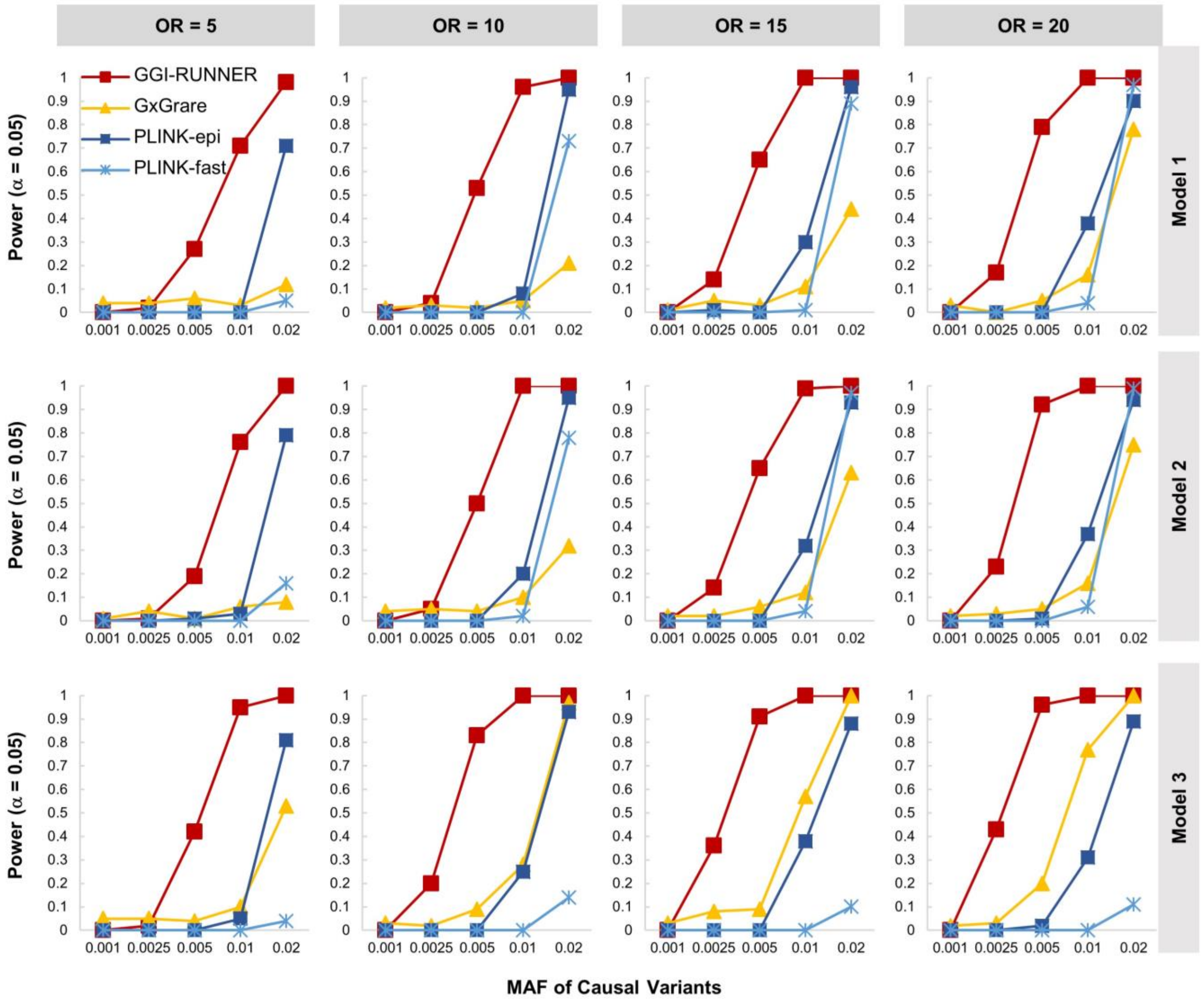


Figure S5. Power curves of four methods: GGI-RUNNER, GxGrare, and two pairwise interaction tests implemented in PLINK, '--epistasis' (PLINK-epi) and '--fast-epistasis' (PLINK-fast). The panels from left to right represent scenarios where the genotype odds ratio (OR) for the causal variants are 5, 10, 15 and 20 with a fixed sample size of 2000 cases and 2000 controls. In each plot, the power of the four methods was tested for assumed causal variants with minor allele frequencies (MAFs) at 0.1%, 0.25%, 0.5%, 1% and 2%. Power was estimated as the proportion of *P*-values less than 0.05 among 100 replicates. Model 1: threshold model, Model 2: multiplicative model, and Model 3: classic epistasis model.

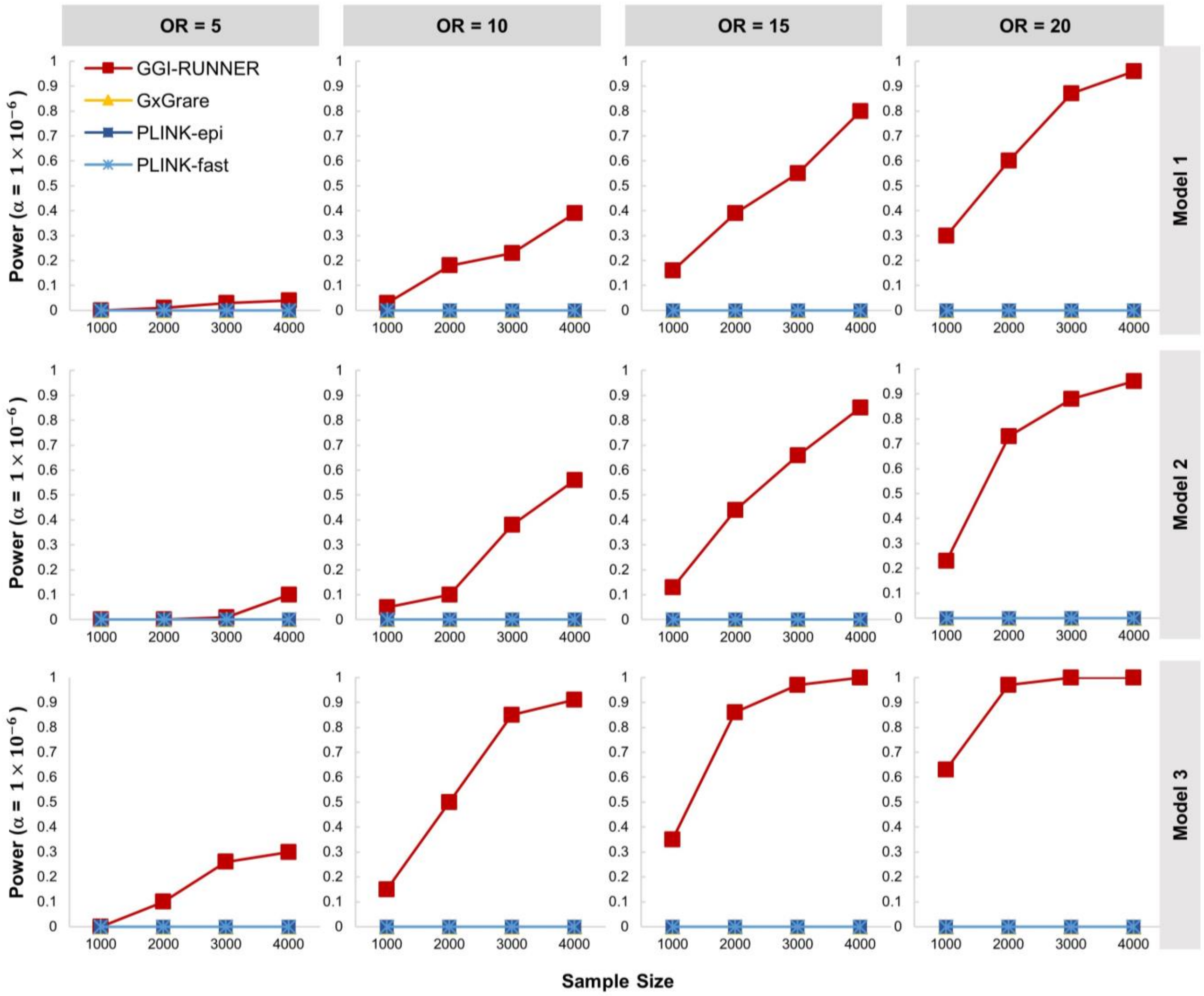


Figure S6. Power curves of four methods: GGI-RUNNER, GxGrare, and two pairwise interaction tests implemented in PLINK, '--epistasis' (PLINK-epi) and '--fast-epistasis' (PLINK-fast). Panels from left to right represent scenarios where the genotype odds ratio (OR) for causal variants are 5, 10, 15 and 20, with a fixed disease prevalence of 0.01 and a minor allele frequency (MAF) of 1%. In each plot, the power of the four methods was tested at sample sizes of 1000, 2000, 3000 and 4000 under a balanced case-control design. Power was estimated as the proportion of $P$-values less than $1 \times 10^{-6}$ among 100 replicates. Model 1: threshold model, Model 2: multiplicative model, and Model 3: classic epistasis model.

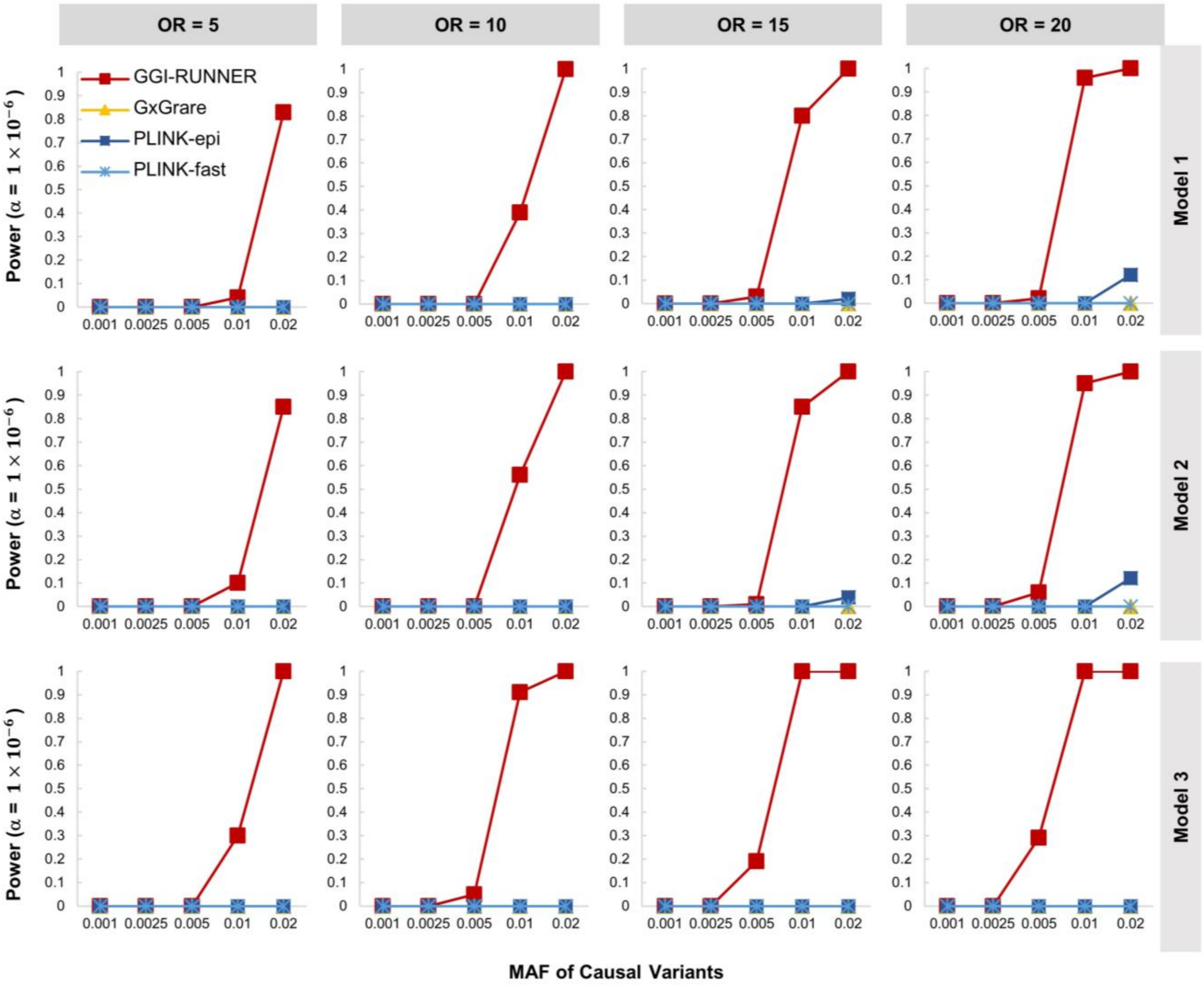


Figure S7. Power curves of four methods: GGI-RUNNER, GxGrare, and two pairwise interaction tests implemented in PLINK, '--epistasis' (PLINK-epi) and '--fast-epistasis' (PLINK-fast). The panels from left to right represent scenarios where the genotype odds ratio (OR) for the causal variants are 5, 10, 15 and 20 with a fixed sample size of 2000 cases and 2000 controls. In each plot, the power of the four methods was tested for assumed causal variants with minor allele frequencies (MAFs) at 0.1%, 0.25%, 0.5%, 1% and 2%. Power was estimated as the proportion of $P$-values less than $1 \times 10^{-6}$ among 100 replicates. Model 1: threshold model, Model 2: multiplicative model, and Model 3: classic epistasis model.

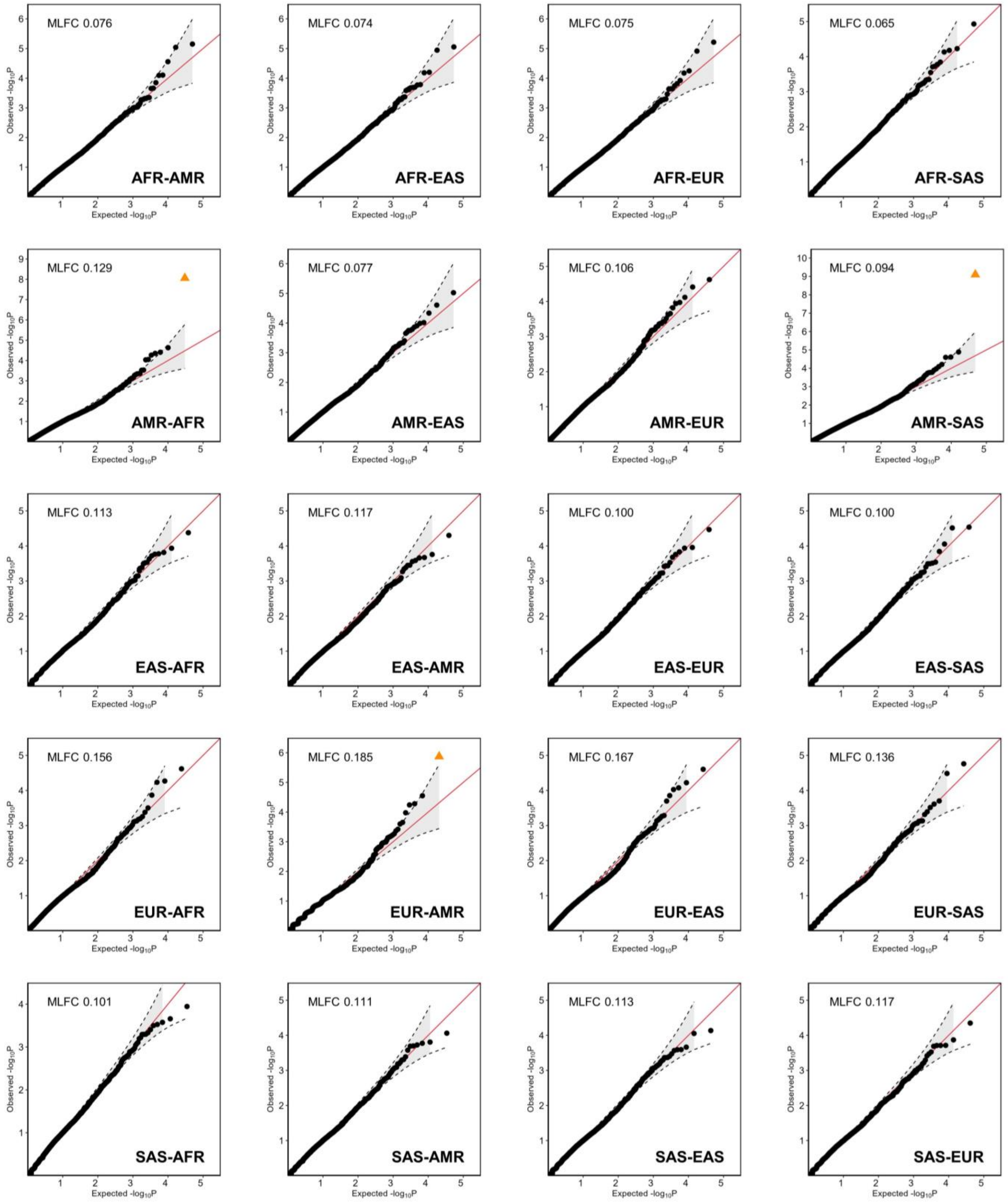


Figure S8. The QQ plots of *P*-values generated by GGI-RUNNER using MAF from ancestral mixed controls. The 20 samples were simulated based on five population panels of the 1KGP. The five panels are AFR- African, AMR- American, EAS- East Asian, EUR- European, and SAS- South Asian. AFR-AMR in the table denotes that the pseudo-cases in this sample were from the AFR panel, while half of the controls were from the AFR panel, and the other half were from the AMR panel. Same as the other 19 group labels. In these analyses, the MAFs from the reference panel of gnomAD matched to the racial background of the cases (e.g., African/African American (AFR) panel of gnomAD for AFR-AMR) are used for rare variant selection. The MAFs from the ancestral mixed controls are used for the accumulated MAF calculation. MLFC: the mean log fold change.

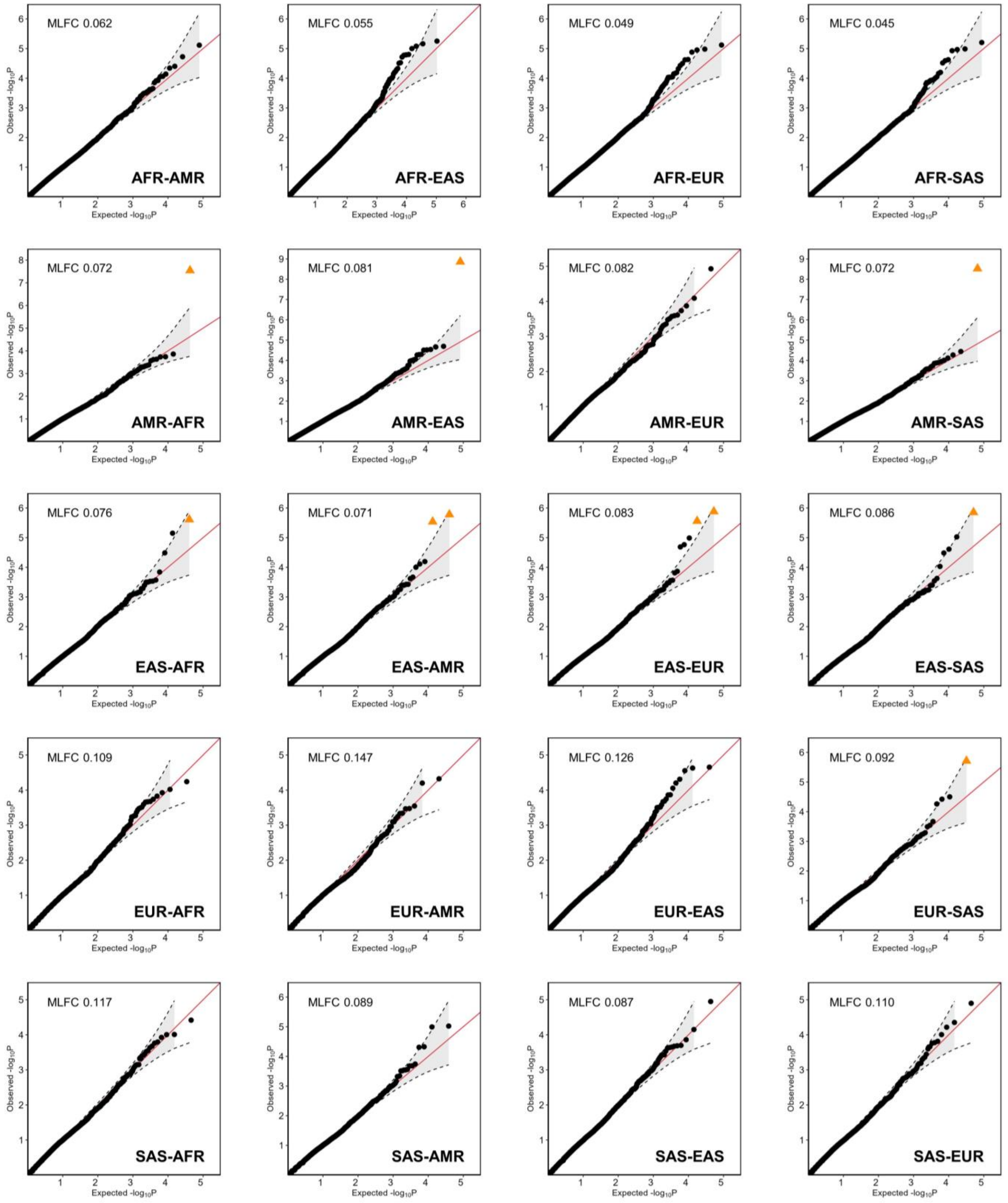


Figure S9. The QQ plots of *P*-values generated by GGI-RUNNER using MAF from ancestry-unmatched reference. The 20 samples were simulated based on five population panels of the 1KGP. The five panels are AFR- African, AMR- American, EAS- East Asian, EUR- European, and SAS- South Asian. AFR-AMR in the table denotes that the pseudo-cases in this sample were from the AFR panel, while half of the controls were from the AFR panel, and the other half were from the AMR panel. Same as the other 19 group labels. In these analyses, the MAFs from the reference panel of gnomAD matched to the racial background of the half ancestral mixed controls (e.g., Latino (AMR) panel of gnomAD for AFR-AMR) are used for rare variant selection and the accumulated MAF calculation. MLFC, the mean log fold change.
